# Targeting DDR1 and DDR2 overcomes matrix-mediated melanoma cell adaptation to BRAF-targeted therapy

**DOI:** 10.1101/857896

**Authors:** Ilona Berestjuk, Margaux Lecacheur, Serena Diazzi, Christopher Rovera, Virginie Prod’homme, Aude Mallavialle, Frédéric Larbret, Sabrina Pisano, Stéphane Audebert, Thierry Passeron, Cédric Gaggioli, Christophe A. Girard, Marcel Deckert, Sophie Tartare-Deckert

## Abstract

Resistance to BRAF and MEK inhibitors in BRAF^V600E^ mutant melanomas remains a major obstacle that limits patient benefit. Microenvironment components including the extracellular matrix (ECM) can support tumor cell adaptation and tolerance to targeted therapies, however the underlying mechanisms remain poorly understood. Here, we investigated the process of matrix-mediated drug resistance (MM-DR) in response to BRAF inhibition in melanoma. We demonstrate that physical and structural cues from fibroblast-derived ECM abrogate anti-proliferative responses to BRAF/MEK inhibition. MM-DR is mediated by the drug-induced clustering of DDR1 and DDR2, two tyrosine kinase collagen receptors. Genetic depletion and pharmacological inhibition of DDR1 and DDR2 overcome ECM-mediated resistance to BRAF inhibition. In melanoma xenografts, targeting DDRs by Imatinib enhances BRAF inhibitor efficacy, counteracts drug-induced collagen remodeling and delays tumor relapse. Mechanistically, DDR-mediated MM-DR fosters a targetable pro-survival NIK/IKKα/NF-κB2 pathway. Our study reveals a novel role of collagen-rich matrix and DDRs in tumor cell adaptation and therapy resistance, thus providing important insights into environment-mediated drug resistance and a pre-clinical rationale for targeting DDR1/2 signaling in combination with BRAF-targeted therapy in melanoma.

## Introduction

One of the hallmarks of cancer cells is their remarkable ability to adapt to microenvironmental influences such as the nature of the stroma, including the extracellular matrix (ECM) and therapeutic stress (Pickup et al., 2014). This is particularly true for malignant skin melanoma, which is one of the most aggressive human cancers and refractory diseases (Shain & Bastian, 2016a). Approximately 50% of skin melanomas carry activating mutations in the *BRAF* oncogene, leading to the activation of the mitogen-activated protein kinase (MAPK)/ERK pathway. Inhibition of the BRAF^V600E/K^ oncoprotein by BRAF inhibitors (BRAFi) such as Vemurafenib has markedly improved clinical outcome of patients (Flaherty et al., 2012). Despite this, durable responses are rare, and most patients relapse within a year of beginning the treatment. Significant prolonged benefit can be achieved by combining BRAFi and MEK (MAPK/ERK kinase) inhibitors (MEKi) such as Trametinib, yet the development of drug resistance remains the most common clinical outcome (Robert et al., 2019). Acquired resistance to targeted drugs involves mostly genetic alterations of key intracellular regulators of the MAPK signaling pathway, leading to restoration of the pathway and non-genetic alterations that are commonly associated with transcriptional reprogramming and phenotype switching from a “proliferative” to an “invasive” cell state characterized by low expression of MITF and SOX10 and high levels of AXL (Rambow et al., 2019). Such adaptive responses to BRAF pathway inhibition are thought to precede mutation-driven acquired resistance (Smith et al., 2016).

However, in addition to mechanisms of resistance intrinsic to the cancer cell, dynamic, *de novo* mechanisms exist, which are orchestrated by the tumor microenvironment and occur during cell adaptation to therapy. Environment-mediated drug resistance (EM-DR) thus appears as an important contributor of cancer cell escape to therapies (Meads et al., 2009). This process has initially been described in multiple myeloma and other hematopoietic malignancies and has been related to minimal residual disease. This phenomenon is gaining importance in the field of melanoma with several studies reporting the involvement of stroma-derived factors in adaptive response and resistance to targeted therapies (Fedorenko et al., 2015, Hirata et al., 2015, Kaur et al., 2016, Straussman et al., 2012, Young et al., 2017). Given the key role of EM-DR enabling the emergence of genetic resistance, an understanding and further identification of EM-DR mechanisms in melanoma may help to develop more effective therapeutic strategies, thereby increasing the efficacy of targeted therapies.

Tumors are complex and adaptive ecosystems that are affected by numerous stromal components that enhance tumor phenotypes and therapy resistance. Cancer-associated fibroblasts (CAFs) are activated fibroblasts and the primary producers of ECM, a highly dynamic structural framework of macromolecules, providing both biochemical and biomechanical cues required for tumor progression (Kalluri, 2016). The ECM is primarily composed of fibrillar and non-fibrillar collagens, hyaluronic acid, proteoglycans, and adhesive glycoproteins such as fibronectin, thrombospondins and SPARC. The ECM also contains matrix-remodeling enzymes and other ECM-associated proteins and acts as a reservoir for cytokines and growth factors. Interactions between cells and ECM elicit intracellular signaling pathways and regulate gene transcription, mainly through cell-surface adhesion receptors including integrins and discoidin domain receptors (DDRs). DDR1 and DDR2 constitute of a unique subfamily of receptor tyrosine kinases and have been identified as non-integrin collagen receptors (Leitinger, 2014, Shrivastava et al., 1997, Vogel et al., 1997). DDR1 and DDR2 are differentiated from each other by their relative affinity for different types of collagens. DDR1 is activated by both fibrillar and non-fibrillar collagens, whereas DDR2 is activated only by fibrillar collagens. Furthermore, their expression and function is associated with fibrotic diseases and cancers. DDR1 or DDR2 are known to control tumor cell proliferation, and invasion, depending on the tumor type and the nature of the microenvironment (Valiathan et al., 2012). However, the functional role of DDR1 or DDR2 kinase activity in mediating sensitivity to anti-cancer therapies and tumor resistance is poorly documented.

Adhesion of tumor cells to ECM is one of the components of EM-DR. However, the influence of matrix-mediated drug resistance (MM-DR) in response to targeted therapies and the nature of ECM receptors driving the MM-DR phenotype in melanoma have not been addressed in detail. To model the contribution of the ECM in melanoma cell responses to BRAF and MEK inhibition, we generated fibroblast-derived 3D ECM from fibroblasts isolated from patient-derived biopsies (melanoma-associated fibroblasts, MAF) and analyzed the process of MM-DR with the aim to identify novel opportunities for microenvironment-targeted therapies. We show that through the pro-survival non-canonical NF-κB2 pathway DDR1 and DDR2 are key mediators of MM-DR in melanoma. Our findings reveal a previously unidentified role of these collagen-activated tyrosine kinase receptors in mediating BRAF inhibitor tolerance and support the rationale to inhibit DDR1 and DDR2 signaling, which disrupts the therapy-resisting properties of the matrix microenvironment. The use of DDR1/2 inhibitors in a novel combinatorial therapeutic strategy may be beneficial for melanoma patients in overcoming resistance to BRAF-targeted therapy.

## Results

### Fibroblast-derived 3D ECM confers drug-protective action to melanoma cells against anti-BRAF^V600E^ therapies

To investigate the potential contribution of MM-DR to targeted therapies in BRAF-mutated melanoma cells, we employed an *in vitro* model of de-cellularized cell-derived 3D ECMs to generate native-like matrices that mimic many structural and biomolecular features typically found *in vivo* (Cukierma et al., 2001). We selected human primary fibroblasts obtained from healthy individuals or purified from patient metastatic melanoma biopsies (melanoma associated fibroblasts, MAFs) and originating from the skin or lymph nodes (LN). The different fibroblast cultures were functionally tested in a 3D collagen matrix contraction assay, which showed that unlike human dermal fibroblasts (HDF), skin and LN MAFs displayed actomyosin contractile activity. Similarly to MAFS, LN normal fibroblasts known as fibroblastic reticular cells (FRC) are myofibroblast-like cells (Fletcher et al., 2015) that showed high propensity to contract collagen (Fig 1A). Cell-derived matrices were then generated denuded of cells and their composition, architecture and rigidity were analyzed using proteomic and microscopic approaches. In our experimental conditions, compared to HDFs, MAFs and FRCs produced and assembled a dense 3D ECM composed of oriented collagen and fibronectin fibers, as shown by picrosirus red and immunofluorescence staining of the ECMs (Fig 1B). Proteomic analysis of the different fibroblast-derived ECMs further documented the molecular composition of these matrices, showing enrichment for several types of collagens (Fig. 1C) and core matrisome components including glycoproteins, proteoglycans as well as ECM regulators and ECM-associated proteins (Supplementary Fig S1). Atomic force microscopy (AFM) analysis of ECM stiffness, revealed values for MAF and HDF matrices that were within range of previous observations (Kaukonen et al., 2016) (Fig 1D). We noticed that matrices generated from FRCs and MAFs were stiffer than HDF-derived ECM. Collectively, these observations validate the use of our experimentally derived matrices for functional studies. Next, we tested the effectiveness of the different fibroblast-derived ECMs to protect BRAF^V600E^-mutated melanoma cells against the anti-proliferative effect of BRAF oncogenic pathway inhibition. We therefore developed a drug protection assay based on the culture of melanoma cells stably expressing a fluorescent nuclear label cultured on fibroblast-derived matrices (Fig 2A). Tumor cells were then treated with different drugs targeting the mutant BRAF/MAPK pathway (Fig 2B). Cell proliferation was monitored using live cell time-lapse imaging and quantified by counting the number of fluorescent nuclei. Cell growth inhibition induced by BRAFi was abrogated when 501MEL melanoma cells were cultured on MAF-derived ECM, in sharp contrast to standard cell culture conditions where cells were plated either on plastic or on purified collagen I (Fig 2C). Thus, MM-DR relies on the topological and molecular features of the ECM. In line with the organization of the 3D matrices depicted in Fig 1, drug protection assays against Vemurafenib revealed that MAF- and FRC-derived matrices display higher protective abilities compared to HDF-derived ECM conferred protection (Fig 2D). Note that throughout the rest of this study we used either FRC- or MAF-derived matrices that exhibit similar functional drug protective activity. Importantly, FRC-derived ECM competently protected 1205Lu cells from the combination of Vemurafenib and Trametinib, which represents the current standard treatment for mutant BRAF melanoma cells (Fig 2E). Cell cycle analysis further showed that experimentally produced ECM from FRCs prevented the G0/G1 cell cycle arrest, induced by the targeted therapy in contrast to the cell culture conditions on plastic (Fig 2F). The ECM therefore transmits signals that prevent the cytostatic action of MAPK pathway inhibitors. At the molecular level, ECM-mediated therapeutic escape of 1205Lu cells from BRAF inhibition was associated with sustained levels of the proliferation markers phosphorylated Rb and E2F1, the survival protein Survivin, and low levels of the cell cycle inhibitor p27KIP1, together with decreased phosphorylation of MEK1/2 and ERK1/2 in presence of the drugs (Fig 2G). Matrices generated by MAFs in addition significantly prevented the inhibitory effect of the dual RAF/MEK inhibitor RO5126766, compared to 501MEL cells on plastic conditions, depicted by the maintained levels of phosphorylated Rb and Survivin and low levels of p27KIP1 (Fig 2H). Upregulation of the pro-apoptotic protein BIM observed in response to RO5126766 was also abrogated in cells plated on fibroblast-derived ECM. Notably, levels of BAX, another pro-apoptotic protein, remained unchanged in the different experimental settings. Similar protective effects were observed when short-term cultures of melanoma cells were plated on FRC- or MAF-derived ECM in presence of Vemurafenib (Fig 2I). Together, these results indicate that fibroblasts assembled and remodeled matrices that provide a drug tolerant environment for BRAF mutant melanoma cell lines and short-term cultures.

**Figure 1.**
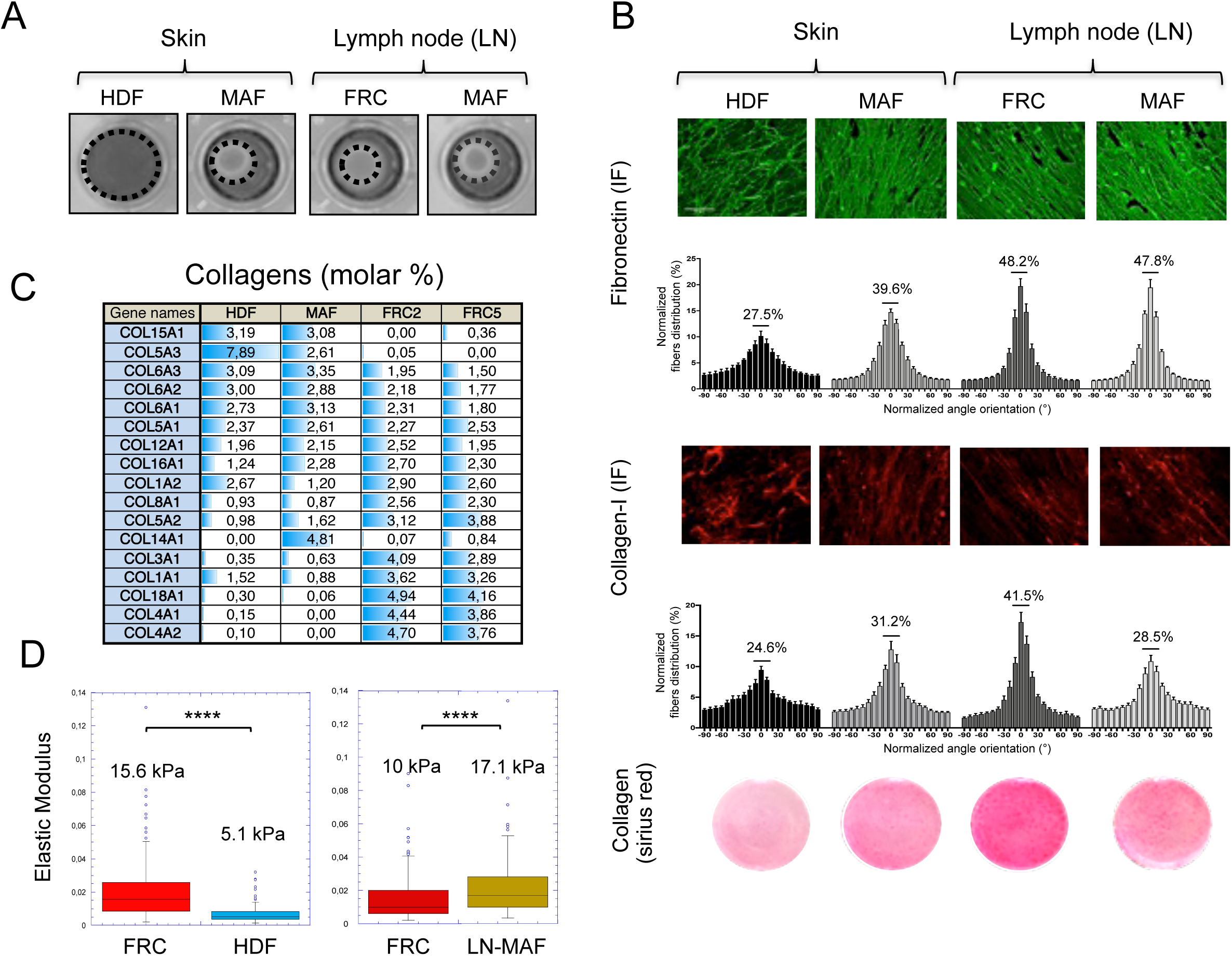
Composition, topology and mechanical properties of fibroblast-derived 3D ECMs. **A** Images show collagen matrix gel contraction by HDF (human dermal fibroblasts), skin-MAF (melanoma associated fibroblasts isolated from skin lesions), LN-FRC (lymph node fibroblast reticular cells), and LN-MAF (melanoma associated fibroblasts isolated from metastatic lymph node). **B** Immunofluorescence analysis of fibronectin (green) and collagen (red) fibers on de-cellularized ECM produced by human fibroblasts. Fibers orientation was quantified using ImageJ software. Scale bar, 10 µm. Images of picrosirius red stained collagen are shown. **C** Collagens identified by mass spectrometry in fibroblast-derived ECM produced by HDF, skin MAF or by two FRCs (FRC#2 and FRC#5). See Supplementary data for the complete list of matrisome core proteins and matrisome-associated proteins (n = 3). **D** Atomic force microscopy (AFM) measurement of the elastic properties (Young’s modulus) of fibroblast-derived ECMs (n = 2).

**Figure 2.**
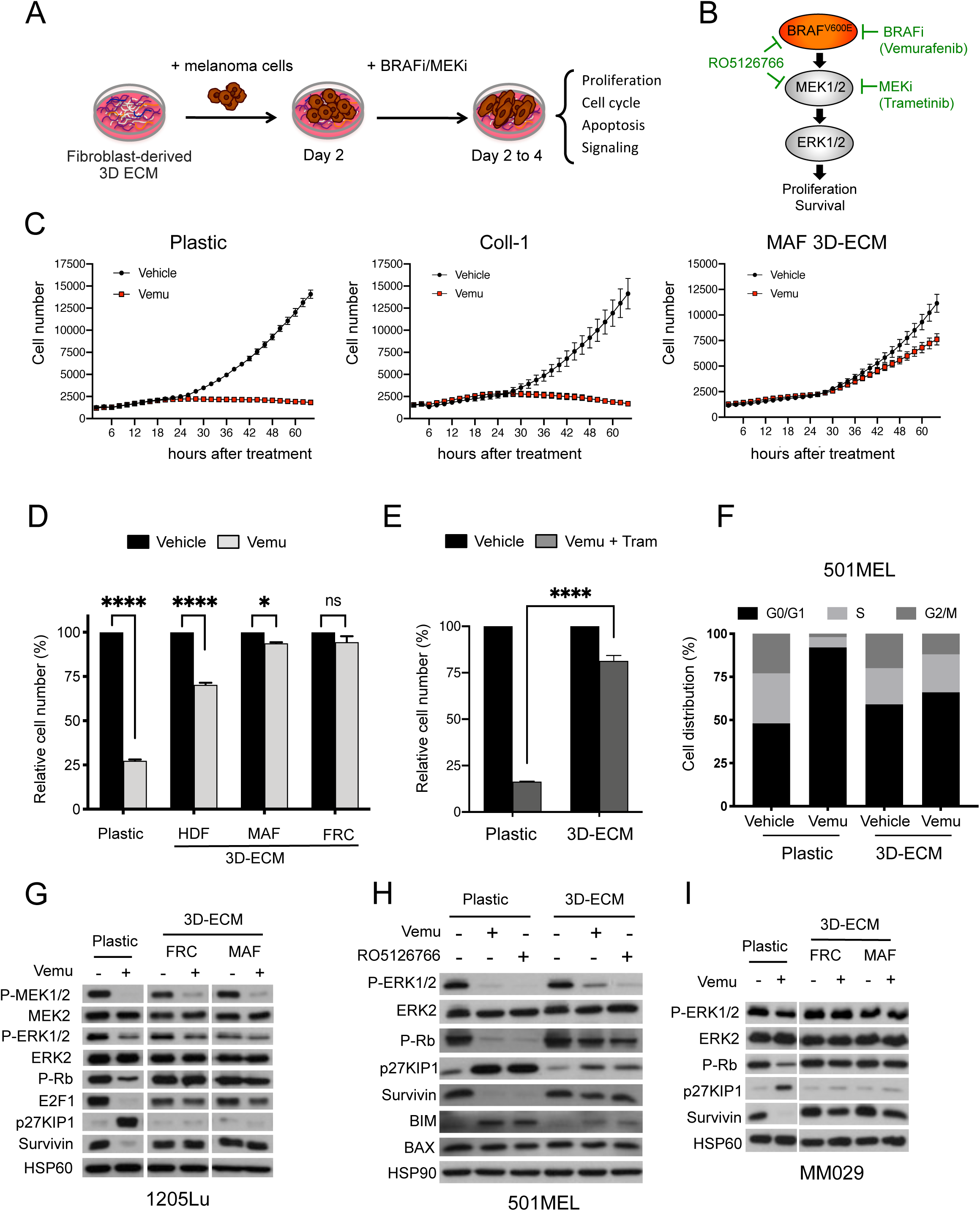
Fibroblast-derived 3D ECM confers drug-protective action to human melanoma cells against anti-BRAF^V600E^ therapies. **A** Scheme of the ECM-mediated drug protection assay. **B** Illustration of the BRAF^V600E^ pathway and the MAPK pathway inhibitors used in the study. **C** Time-lapse imaging of the proliferation of NucLight-labeled 501MEL cells plated on plastic (left panel), Coll-1 (collagen-1; middle panel) or LN-MAF-derived ECM (right panel) treated with vehicle or 2 µM Vemurafenib using the IncuCyte ZOOM system. **D** Quantification of proliferation of NucLight-labeled 1205Lu cells plated for 48 h on plastic or on the indicated fibroblast-derived ECM prior a 96 h treatment with 5 µM Vemurafenib. *P<0.05, ****P<0.001, two-way ANOVA followed by Sidak’s multiple comparison test. **E** Quantification of the proliferation of 1205Lu cells plated for 48 h on plastic or FRC-derived-ECM prior treatment with 2 µM Vemurafenib and 0.01 µM Trametinib for 96 h. ****P<0.001, two-way ANOVA followed by Sidak’s multiple comparison test. **F** Flow cytometry analysis of cell cycle distribution of 501MEL cells cultured on plastic or MAF-derived ECM and treated with vehicle or 2 µM Vemurafenib. The percentage of cells in different phases of the cell cycle is indicated. **G, H, I** Immunoblotting of protein extracts from 1205Lu and 501MEL cell lines and short-term melanoma cultures (MM029) cultivated as described above on plastic or the indicated fibroblast-derived ECM in the presence or not of Vemurafenib 5 µM (G), 2 µM (H, I), or the dual RAF/MEKi RO5126766 1 µM (H) for 48 h, using antibodies against P-MEK1/2, P-ERK1/2, P-Rb, E2F1, Survivin, p27KIP1, BIM or BAX. HSP60, loading control.

### Expression of the collagen receptors DDR1 and DDR2 in melanoma

Previous studies have demonstrated the critical role of ECM receptors belonging to the integrin family in drug resistance (Seguin et al., 2015). Moreover, BRAF inhibition has been described to generate a drug protective mesenchymal stroma with high β1 integrin/FAK signaling, as a result of the paradoxical action of BRAFi on MAFs (Hirata et al., 2015). Yet, in our experimental settings, we were unable to demonstrate a significant implication of the β1 integrin /FAK axis in drug protection conferred by fibroblast-derived ECMs. Indeed, addition of blocking β1 integrin antibodies or depletion of FAK both failed to prevent the protective property of fibroblast-derived ECM against the growth and survival inhibitory signals induced by BRAFi (Supplementary Fig S2).

This prompted us to interrogate the contribution of other ECM receptors in drug tolerance. Keeping in mind the elevated levels of fibrillar collagens found in fibroblast matrices (Fig 1C), we examined the functional implication of the collagen tyrosine kinase receptors DDR1 and DDR2 (Shrivastava et al., 1997, Vogel et al., 1997). The analysis of TCGA datasets for cutaneous melanoma showed that *DDR1* and *DDR2* genes were genetically altered in 20% and 13% of melanoma cases, respectively. Interestingly, a significant fraction of melanomas was found to be associated with high mRNA levels of *DDR1* and *DDR2* (respectively 11% and 7%), consistent with the notion that these collagen receptors may play an important role in the pathogenicity of melanoma (Fig 3A). Immunohistochemical analysis of DDR1 and DDR2 expression in benign (naevus) and malignant melanocytic skin lesions (primary and metastatic), further showed that DDR1 and DDR2 levels significantly increased during melanoma progression, indicating that DDR1 and DDR2 may represent novel prognostic factors in melanoma (Fig 3B). Tumor cell plasticity is critical during melanoma progression and therapeutic response (Arozarena & Wellbrock, 2019). We thus examined the levels of DDR1 and DDR2 in a collection of melanoma cell lines and short-term melanoma cultures, according to cell state differentiation markers. Both DDR1 and DDR2 were expressed in “classic” melanoma cell lines. Notably, high levels of DDR2 were found associated with low levels of melanocytic markers MITF and SOX10 and high levels of the invasive marker AXL in metastatic 1205Lu cells (Fig 3C) and in BRAFi-resistant M229R and M238R cells (Fig 3D). In short-term melanoma cultures, both DDR1 and DDR2 were detected in MM099 and MM029 cells with the MITF^low^, SOX10^low^ and AXL^high^ invasive phenotype signature (Fig 3E). The examination of additional public gene expression datasets confirmed that *DDR2* levels were significantly linked to an AXL^high^ melanoma invasive cell population across the Mannheim, Philadelphia and Zurich cohorts (GSE4843, GSE4841 and GSE4840), whereas *DDR1* expression was similarly detected both in MITF^low^ proliferative and AXL^high^ invasive cell phenotypes (Fig 3F) (Widmer et al., 2012). Together, these observations associate DDR1 and DDR2 expression with melanoma progression and link DDR2 to the invasive and therapy-resistant melanoma cell sub-population.

**Figure 3.**
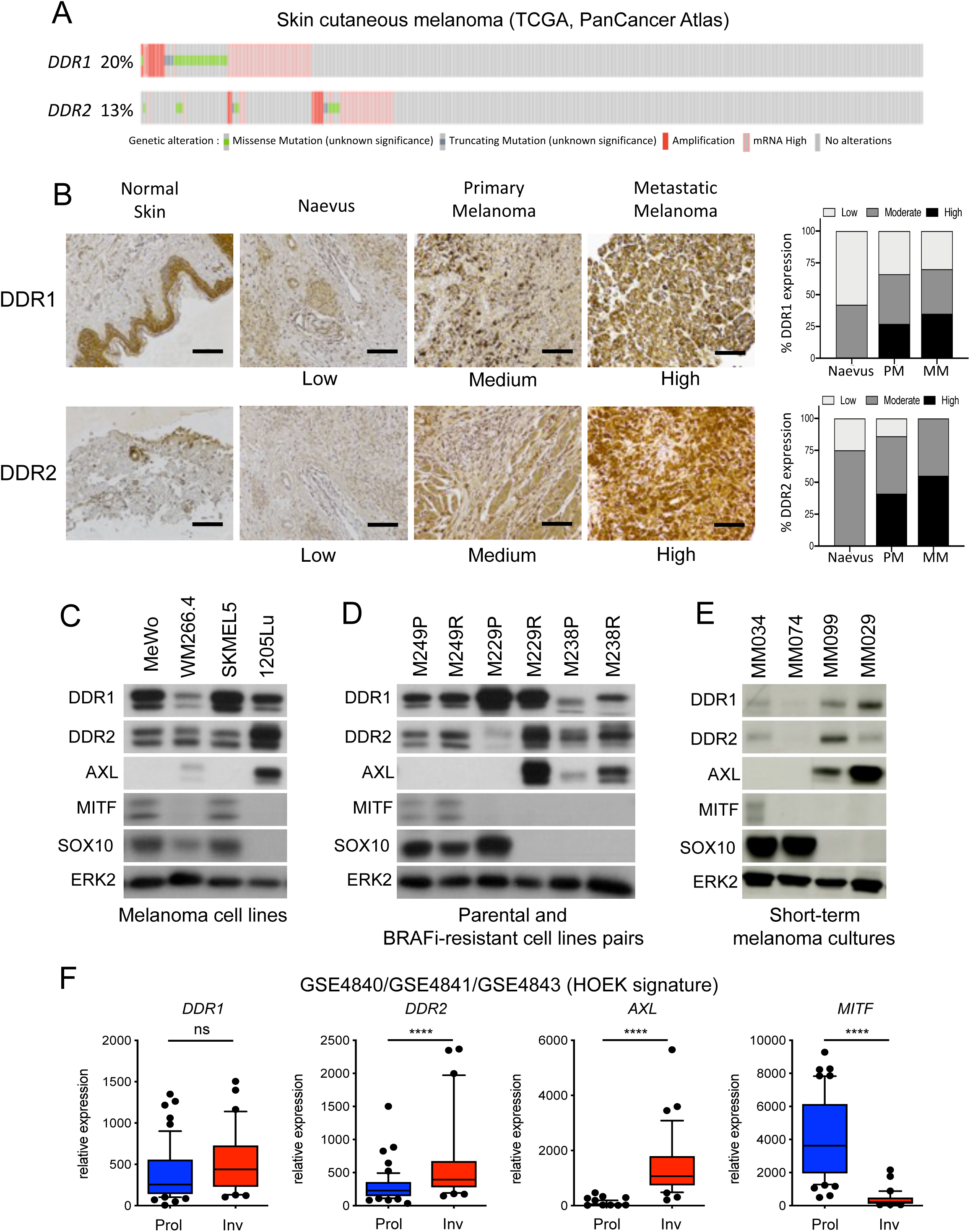
Expression of DDR1 and DDR2 in human melanoma. **A** Meta-analysis of 363 cutaneous melanoma from TCGA (skin cutaneous melanoma, PanCancer Atlas) (http://www.cbioportal.org/) showing the percentage of samples with genetic alterations of *DDR1* and *DDR2*. Cases with missense (green) and truncating (blue) mutations, amplification (red) and mRNA overexpression (pink) are indicated; gray, individual cases. **B** Immunohistochemical analysis of DDR1 and DDR2 levels on human melanoma tissue microarrays. Representative IHC images and quantification (right bar histograms) of DDR1 and DDR2 expression in normal skin, naevus, primary melanoma (PM) and lymph node melanoma metastases (MM). Scale bar, 100 µm. Samples were scored as low, medium or high for DDR1 or DDR2 expression (naevus, n = 12; PM, n = 30; MM, n = 20). **C, D, E** Immunoblotting of equal amounts of protein extracts from melanoma cell lines (C), isogenic pairs of parental sensitive and BRAFi-resistant cell lines (D) or short-term cell cultures from BRAF mutant patients using antibodies against DDR1, DDR2 or markers of the melanoma cell phenotype switching AXL, MITF or SOX10. ERK2, loading control. **F** Box and whisker plots (10th to 90th percentile) show *DDR1* and *DDR2* expression across proliferative (Prol, n=53) *versus* invasive (Inv, n=33) cell states of melanoma cultures within the Mannheim, Philadelphia, and Zurich cohorts (GSE4840, GSE4841 and GSE4843, respectively data sets). The expression of *AXL* and *MITF* is shown as differentiation control markers. ns, non significant (p>0.05); ****p<0.0001 two-tailed Mann Whitney test.

### Targeting DDR1 and DDR2 impairs ECM-mediated resistance to oncogenic BRAF pathway inhibition

To address the contribution of DDR1 and DDR2 in MM-DR, a siRNA approach was used to target DDR1, DDR2 or both in melanoma cells cultured on fibroblast-derived ECM, in presence of BRAF and/or MEK inhibitors. Immunoblot analysis showed specific DDR1 and DDR2 protein reduction after siRNA transfection in BRAFi-treated 1205Lu cells (Fig 4A) and in BRAFi/MEKi-treated MM099 cells (Fig 4B). Compared to the single knockdown, the simultaneous knockdown of DDR1 and DDR2 overcame MM-DR to BRAF targeted therapy as revealed by decreased levels of phospho-Rb and Survivin in 1205Lu cells (Fig 4A). Similar results were obtained using a second combination of DDR1/2-targeting siRNA sequences (Supplementary Fig S3A). In addition, the depletion of both DDR1 and DDR2 enhanced the cytotoxic activity of BRAF/MEK co-targeting as shown by the increased cleavage of apoptotic Caspase 3 that was detected in MM099 cells (Fig 4B). DDRs are druggable receptors, which are targeted by Imatinib, a tyrosine kinase inhibitor (TKI) developed as an ABL inhibitor but was later shown to inhibit DDR1 and DDR2 kinase activities with high efficacy (Day et al., 2008). Imatinib belongs to therapeutic molecules used in the clinic for the treatment of chronic myeloid leukemia (CML) and acute lymphoblastic leukemia (ALL) with the Philadelphia chromosome (Druker et al., 2001). Enzymatic activities of DDR1 and DDR2 were also inhibited by other small molecules including DDR1-IN-1 (Kim et al., 2013). We first confirmed that Imatinib and DDR1-IN-1 efficiently inhibited type I collagen-induced DDR1 and DDR2 tyrosine phosphorylation in 1205Lu cells (Fig 4C). Inhibition of DDR1/2 kinases by Imatinib or DDR1-IN-1 suppressed the protective action of MAF-derived ECM against Vemurafenib treatment as evidenced by the strong decrease in cell proliferation observed in 1205Lu cells co-treated with Vemurafenib and DDR1/2 inhibitors (Fig 4D). Similar potentializing effects of DDR1/2 inhibitors were observed in drug-protection assays performed on FRC-derived matrices (Supplementary Fig S3B). Mutant BRAF and DDR1/2 co-targeting in melanoma cells plated on MAF-derived ECM decreased levels of phosphorylated Rb and Survivin, and induced Caspase 3 cleavage (Fig 4E). Similar biochemical events were promoted by BRAFi or the combination BRAFi/MEKi in the presence of Nilotinib, a next-generation BCR-ABL inhibitor approved for the treatment of Imatinib-resistant patients and targeting DDR1/2 (Day et al., 2008) (Supplementary Fig S3C). Induction of apoptosis in co-treated melanoma cells was further confirmed using flow cytometry analysis of cell death markers (Fig 4F).

**Figure 4.**
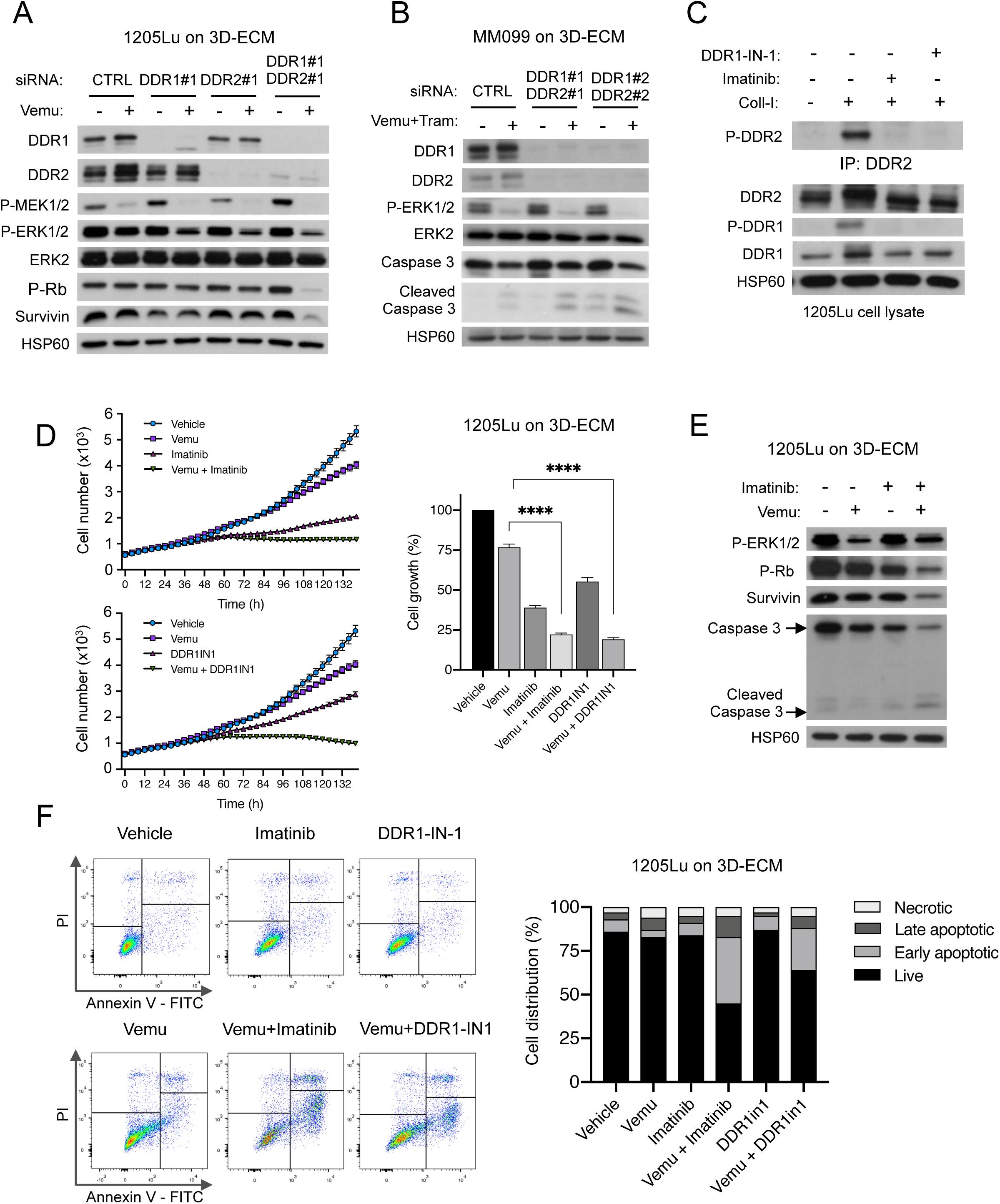
Inhibition of DDR1 and DDR2 by genetic or pharmacological approaches abrogates ECM-mediated resistance to BRAF^V600E^ pathway inhibition. **A** Immunoblotting of protein extracts of siCTRL-, siDDR1#1-, siDDR2#1- or siDDR1#1/siDDR2#1- transfected 1205Lu cells plated on MAF-derived ECM treated with vehicle or 5 µM Vemurafenib for 96 h, using antibodies against DDR1, DDR2, P-MEK1/2, P-ERK1/2, P-Rb or Survivin. HSP60, loading control. **B** Immunoblotting of protein extracts from short-term melanoma cell cultures MM099 transfected with siCTRL-, the combination siDDR1#1/siDDR2#1 or siDDR1#2/siDDR2#2 prior being cultivated on FRC-derived ECM and treated with vehicle or 2 µM Vemurafenib combined with 0.01 µM Trametinib, using antibodies against DDR1, DDR2, P-ERK1/2 or cleaved Caspase 3. HSP60, loading control. **C** Immunoblot analysis of collagen I (Coll-I)-induced DDR1 and DDR2 tyrosine phosphorylation in 1205Lu cells. Cells were incubated with 10 µg/ml of Coll-I in the presence or not of 7 µM Imatinib or 1 µM DDR1-IN-1 for 18 h. After cell lysis, DDR2 autophosphorylation was analyzed with anti-P-DDR2 following immunoprecipitation (IP) with anti-DDR2 antibodies. DDR1 autophosphorylation was analyzed in total cell lysates with anti-P-DDR1. HSP60 was used as loading control. **D** Time-lapse imaging of proliferation of NucLight-labeled 1205Lu cells plated for 48 h on MAF-derived ECM prior treatment with 5 µM Vemurafenib in the presence or not of 7 µM Imatinib or 1 µM DDR1-IN-1 for the indicated time (left). The histograms show quantification of cell proliferation at the experiment end point (right). ****P<0.001, Kruskal-Wallis test followed by Dunn’s multiple comparison test. **E** Immunoblotting of protein extracts from 1205Lu cells cultivated on MAF-derived ECM, treated or not with 5 µM Vemurafenib and/or 7 µM Imatinib using antibodies against P-Rb, P-ERK1/2, Survivin or cleaved Caspase 3. HSP60, loading control. **F** Flow cytometry analysis of cell death (Annexin V/PI labeling) in 1205Lu cells plated on FRC-derived ECM and treated with vehicle or 5 µM Vemurafenib in the presence or not of 7 µM Imatinib or 1 µM DDR1-IN-1. Right bar histograms show the distribution of cells (% of total) across the different forms of death.

Recent studies have described that collagen binding to DDR1 and DDR2 leads to their activation and clustering and into filamentous membrane structures associated with collagen fibrils (Yeung et al., 2019). We thus examined the spatial distribution of DDR1/2 in melanoma cells plated on collagen I coated plastic dishes or on fibroblast-derived ECM. Immunofluorescence staining of phosphorylated DDR1 and DDR2 revealed that cells cultivated on purified collagen I displayed a globular dot-like distribution of the two receptors, whereas cells seeded on 3D ECM exhibited a significant fraction of DRR1/2 distributed into linear membrane clusters (Fig 5A and B). Interestingly, cell exposure to Vemurafenib dramatically increased the proportion of DDR1/2-containing linear clusters on 3D ECM (Fig 5A and B). Concomitantly, melanoma cell spreading was increased upon BRAF inhibition albeit to lesser extent on 3D ECM (Fig 5C). These data suggests that BRAFi-induced cytoskeletal changes drive the clustering of DDRs along collagen fibers and subsequent MM-DR (Fig 5D).

**Figure 5.**
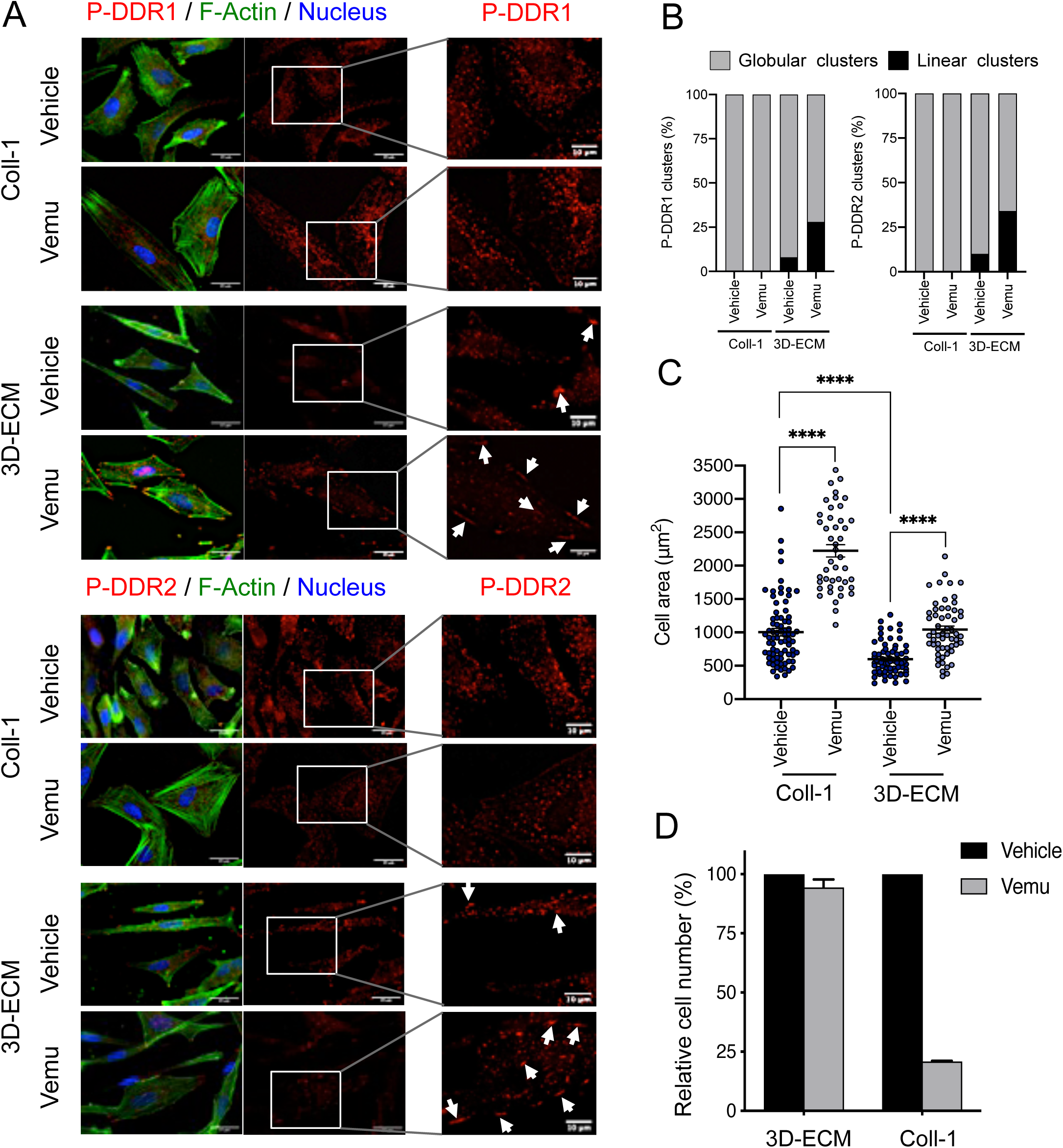
Interaction of melanoma cells with 3D-ECM induces the clustering of DDR1 and DDR2 upon BRAFi treatment. **A** Representative images of 1205 cells cultivated on collagen-I (Coll-I) or FRC-derived ECM for 48 h prior treatment with vehicle or 5 µM Vemurafenib for 96 h. Immunofluorescence for Phospho-DDR1 (P-DDR1) (red; top panels) and Phospho-DDR2 (P-DDR2) (red; bottom panels), F-actin (green) and nuclei (blue) is shown. Enlarged images of P-DDR1 and P-DDR2 immunostaining are shown. White arrows indicate P-DDR1 and P-DDR2 cell membrane linear clustering. Scale bar = 10 µm. **B, C** Quantification of globular *versus* linear clusters of phosphorylated DDR1 and DDR2 (B) and of cell area (C) from immunofluorescence staining shown in (A) using Image J software. Data are from >50 individual cells (n=3). ****P<0.0001, Kruskall-Wallis test followed by Dunn’s multiple comparisons test. **D** Quantification of 1205Lu cell proliferation following cultures on Coll-I or FRC-derived 3D-ECM in the presence or not of 5 µM Vemurafenib showing as a control the drug protective effect of fibroblast-derived ECM against BRAFi compared to Collagen I coating.

Together, these findings suggest that DDR1 and DDR2 determine BRAF mutant melanoma cell responsiveness to targeted therapies and that the drug-tolerance action of DDRs is dependent to their enzymatic activities and plasma membrane distribution.

### Pharmacological inhibition of DDR1/2 by Imatinib improves BRAFi efficacy, counteracts drug-induced collagen remodeling and delays tumor relapse

The anti-tumor activity of Imatinib combined with Vemurafenib was next assessed in a pre-clinical xenograft model of melanoma. BRAF-mutated melanoma cells 1205Lu were subcutaneously xenografted into nude mice (CDX model), which were exposed to Vemurafenib, Imatinib or Vemurafenib plus Imatinib (Fig 6A). As expected, BRAF inhibition induced a rapid tumor reduction, whereas Imatinib alone did not display a significant anti-melanoma effect (Fig 6B). However, after 12 days, tumors treated with BRAFi alone had resumed growth, whereas the combination with Imatinib markedly delayed tumor relapse and led to a significant reduction of tumor volume (Fig 6B and D) and weight (Fig 6C). Immunohistochemical analysis of tumor samples further documented that in comparison to single regimens, the combined treatment with Vemurafenib and Imatinib dramatically reduced cell proliferation and led to apoptotic tumor cell death as shown by *in situ* expression of cleaved Caspase 3 and decreased Ki67 (Fig 6D). Consistently, the Vemurafenib/Imatinib combination significantly increased the survival of melanoma-bearing mice (Fig 6E) treated for 30 days without apparent body weight loss or signs of toxicity throughout the study (Fig 6F). Increased collagen deposition has previously been described in melanoma xenografts upon treatment with Vemurafenib (Jenkins et al., 2015). We next investigated collagen content and fiber organization in tumor tissues from mice treated with the mono- or combo-therapy. Histochemical analysis showed that Vemurafenib treatment triggered a profound remodeling of the melanoma ECM stroma (Fig 7A), with marked increase of the collagen fibers’ area and thickness, which was suppressed by Imatinib. Collagen color analysis under polarized light showed a decrease of mature orange and red fibers in tumors treated with the combined regimen compared to the single-agent treatment (Fig 7B). Finally, tumor imaging with second harmonic generation (SHG) confirmed the fibrillar nature of the collagen network that was rearranged upon Vemurafenib treatment but not upon the combined Vemurafenib/Imatinib treatment (Fig 7A and C). These data suggest that treatment with Imatinib counteracts the adverse effect of targeted agents on aberrant collagen deposition and organization, a process potentially contributing to drug resistance and relapse.

**Figure 6.**
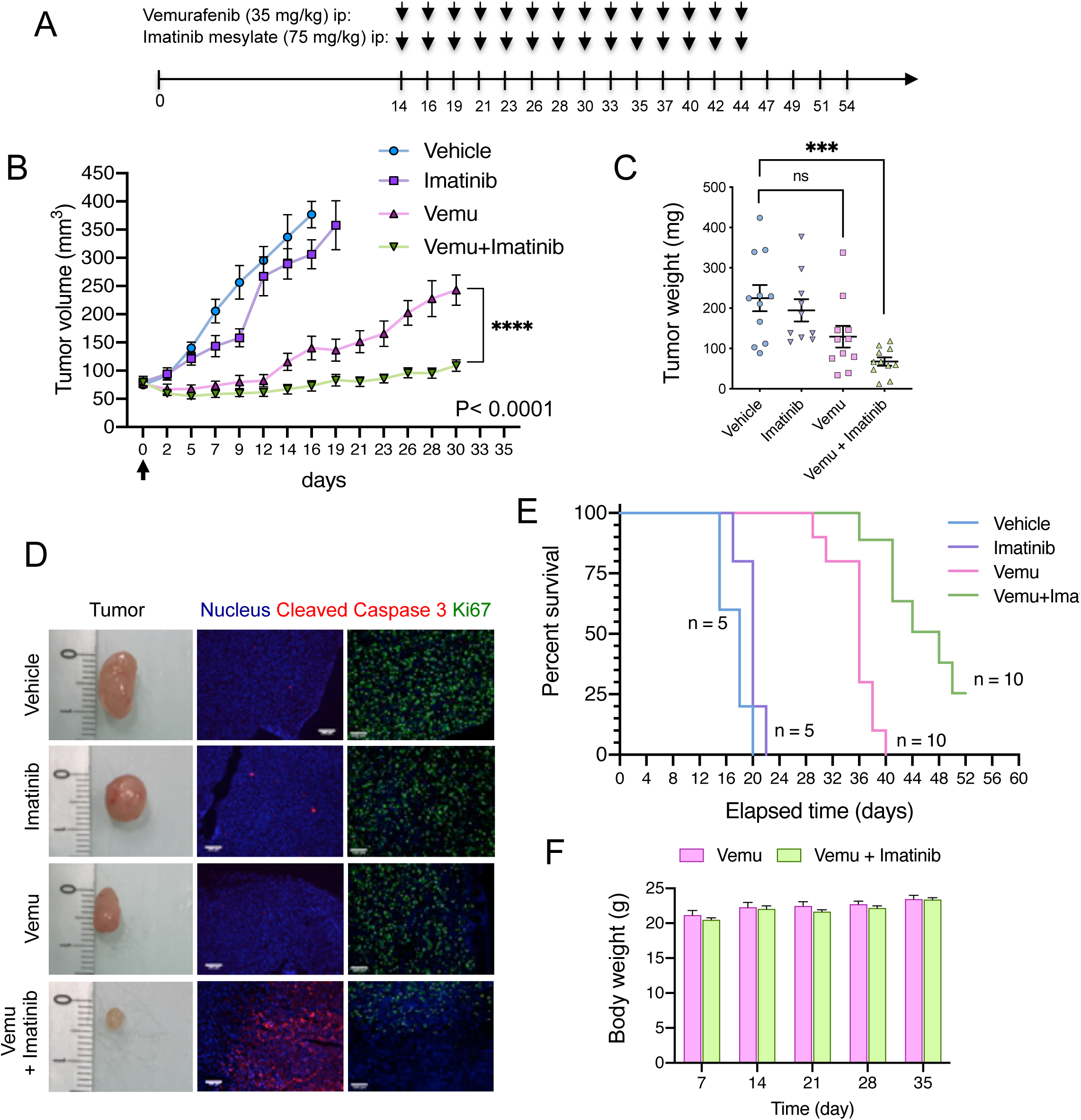
Targeting DDR1 and DDR2 by Imatinib sensitizes melanoma tumors to BRAF^V600E^ inhibition. **A** Outline of the experimental set up and treatment regimens. **B** 1205Lu cells were s.c. inoculated into nude mice and when tumors reached 75 mm^3^, mice were treated with the indicated mono- or combo-therapy for 30 days. Graphs show tumor growth following treatment by indicated drugs. Data shown are mean ± SEM of tumor volume (n = 6; ****P<0.0001, two-way ANOVA followed by Tukey’s multiple comparison test). **C** Scatter plot graphs showing the tumor weight upon treatment by the indicated mono- or combo-therapy. Data shown are mean ± SEM of tumor weight. ***P<0.0002, Kruskall-Wallis test followed by Dunn’s multiple comparison test; ns, non significant. **D** Immunofluorescence stainings using anti-cleaved Caspase 3 (red), anti-Ki67 (green) and DAPI in tumor sections of 1205Lu-derived xenografts from (B). Scale bar, 100 µm. A microphotograph of the tumor size in each treated group is shown (representative of n=5 to 10). **E** Kaplan-Meier survival curves of mice treated with vehicle, Imatinib, Vemurafenib or Vemurafenib/Imatinib. Median time to progression was 18, 20, 36 and 48 days, respectively. Log rank (Mantel-Cox) for Vemurafenib *vs* Vemurafenib/Imatinib mesylate. ****p < 0.0001. **F** Mouse body weight was measured at the indicated day. Data shown are mean ± SEM (n = 10).

**Figure 7.**
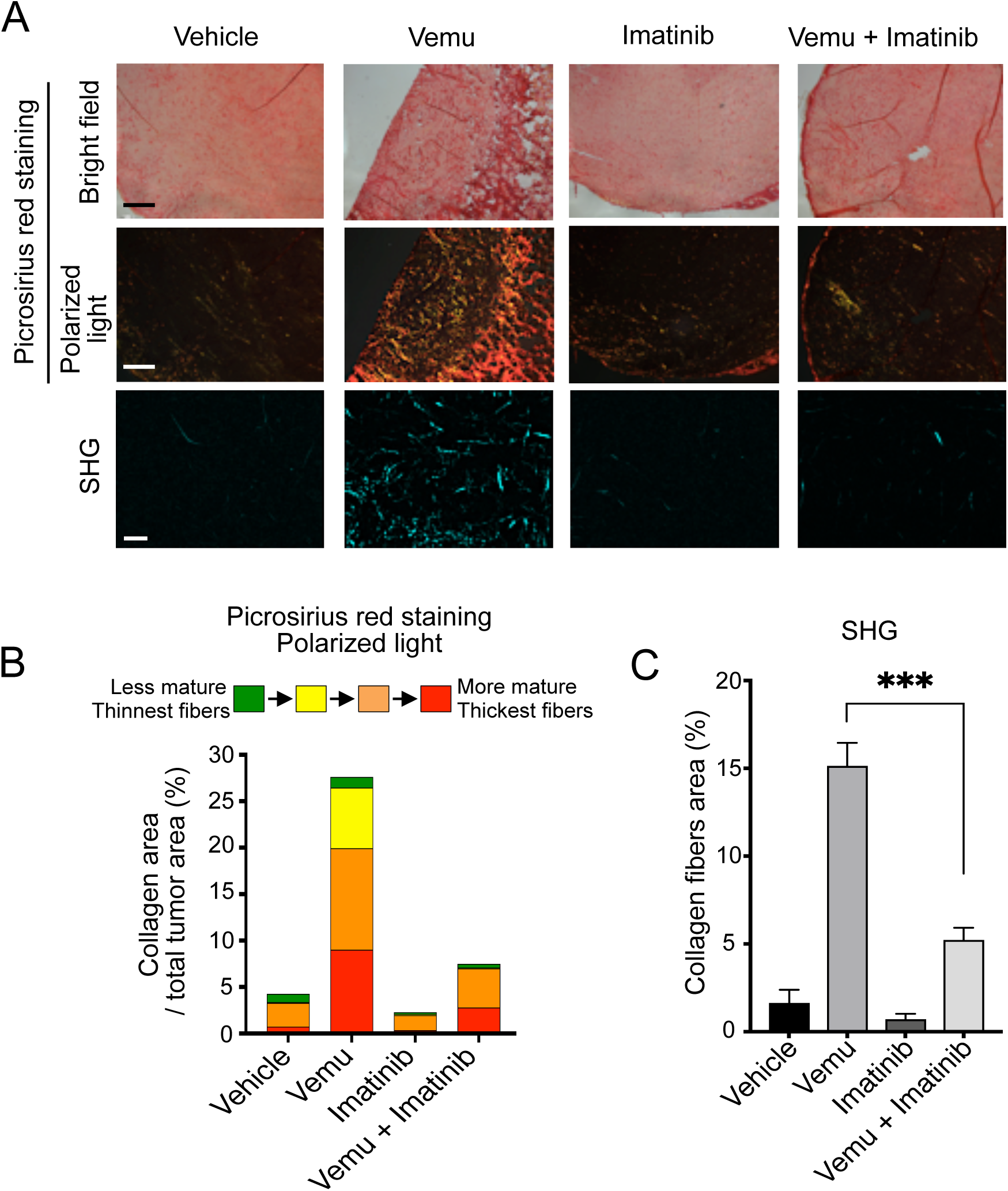
Imatinib normalizes collagen deposition and remodeling induced upon BRAFi treatment. **A** Sections of 1205Lu xenografts from Fig. 6B were stained with picrosirius red and imaged under transmission light (upper panels) or polarized light (middle panels) (scale bar, 500 µm) or imaged by second harmonic generation (SHG) microscopy (lower panels) (scale bar, 50 µm) to examine collagen fiber network upon the mono or combined regimens. **B** Quantification of collagen maturity and fiber thickness in 1205Lu xenografts stained with picrosirius red using polarized light microscopy. Birefringence hue and amount of collagen fibers were quantified as a percent of total tissue area. **C** Quantification of collagen fibers using SHG microscopy in tumor sections from (A). ***p< 0.0002, Mann-Whitney test.

### DDR1/2-mediated MM-DR involves a targetable pro-survival NIK-IKKα-NF**κ**B2 pathway

Finally, we wished to better characterize at the molecular level the MM-DR process that is promoted by collagen receptors DDR1 and DDR2. In order to identify MM-DR-related signaling pathways triggered by their tyrosine kinase activity, we performed phospho-kinase screening with protein extracts from melanoma cells cultured on fibroblast-derived matrices or plastic, in the presence of Vemurafenib. Unfortunately, this approach did not reveal any significantly augmented kinase activity or increased phosphorylation of kinase substrates (Supplementary Fig S4). Moreover, consistent with the results of the phospho-kinase screening, we failed to observe by immunoblot analysis any changes in the activity of the AKT survival pathway in 1205Lu and 501MEL cells cultured under MM-DR conditions (Supplementary Fig S5). On the contrary, biochemical analysis revealed that unlike single-agent treatment, the combined treatment with Vemurafenib and Imatinib dramatically reduced the levels of RelB and phosphorylated p65 in ECM-stimulated melanoma cells following 96 h (Fig 8A). RelA (p65) and RelB are components of the canonical and non-canonical NFκB pathway, respectively (Taniguchi & Karin, 2018). Previous studies have linked the NFκB pathway to melanoma invasion (Rathore, Girard et al., 2019) and growth (De Donatis et al., 2016), as well as intrinsic and environment-mediated drug resistance to MAPK pathway inhibitors (Konieczkowski et al., 2014, Smith et al., 2014, Young et al., 2017). Consistently, siRNA-mediated concomitant depletion of DDR1 and DDR2, which impaired melanoma MM-DR by triggering cell cycle arrest and apoptosis, decreased the expression of RelB, NFκB2 precursor protein p100 and NFκB2 p52 processed form in melanoma cells cultured in MM-DR conditions (Fig 8B). The non-canonical NFκB2/p52-RelB pathway is activated by the upstream kinases NIK (NFκB inducing kinase) and IKKα (IκB kinase) (Fig 8C). Using a pan-IKK inhibitor BMS345541 (Fig 8D) and a recently developed NIK inhibitor (NIKi) (Mondragon et al., 2019) (Fig 8E), we confirmed the implication of the NFκB2 pathway in melanoma MM-DR, as illustrated by decreased cell cycle markers and increased apoptotic markers in BRAFi-treated ECM-stimulated melanoma cells. Similar inhibition of p52 and RelB was found in melanoma cells transfected with DDR1/2-targeting siRNA after exposure to Vemurafenib and Trametinib (Supplementary Fig S6A). Finally, comparable decreased levels of RelB, p100 and p52 were observed following pharmacological targeting of DDR1/2 and NIK in MM-DR conditions (Supplementary Fig S6B). Together, our findings suggest that melanoma cell adaptation to BRAFi/MEKi treatment involves the interaction of tumor cells with the mesenchymal stroma enriched in fibrillar collagen, thereby promoting DDR1/2-dependent activation of the NFκB2-RelB pathway (Fig 9).

**Figure 8.**
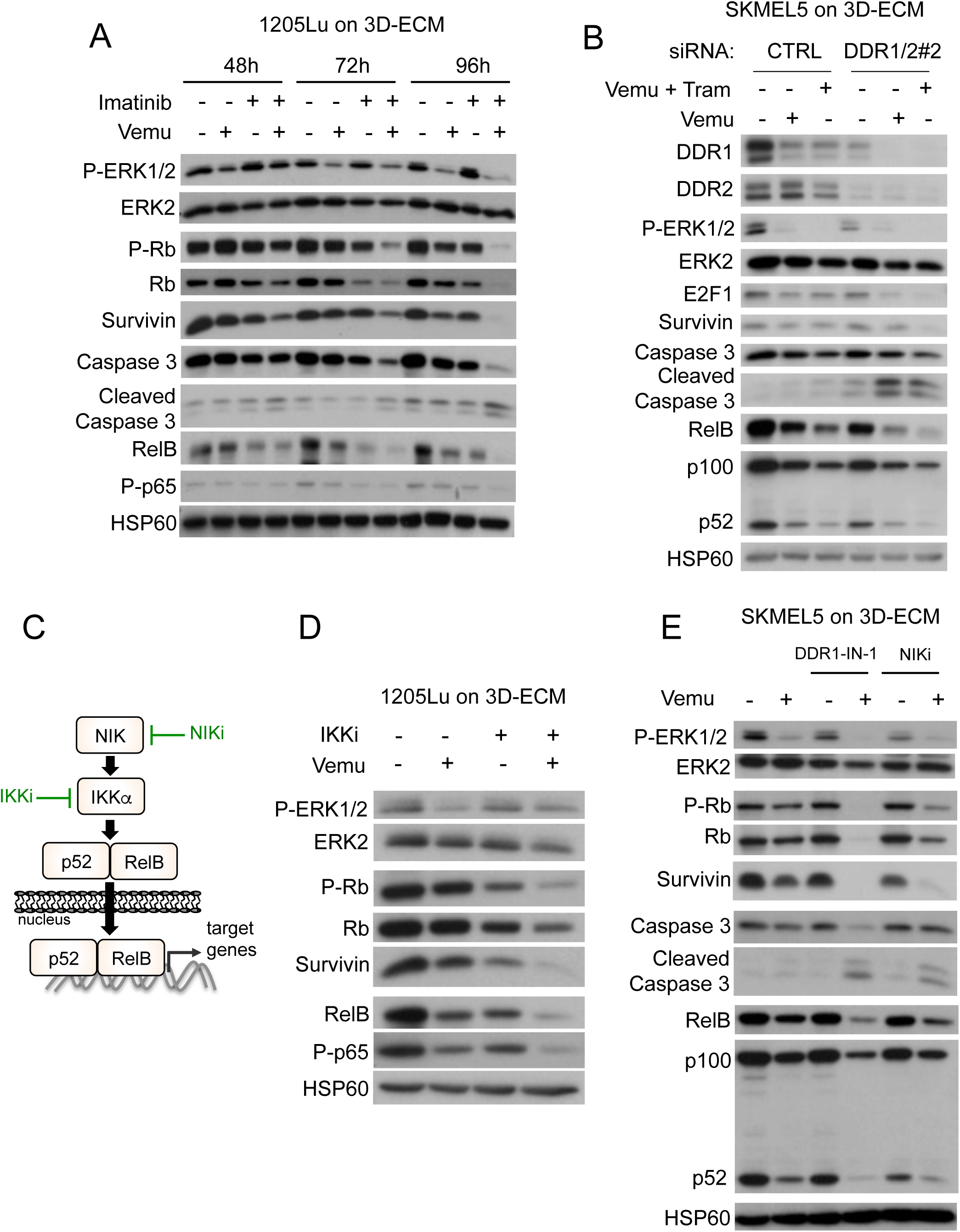
Targeting the non-canonical NFκB2 pathway overcomes DDR1/2-dependent MM-DR to BRAF/MEK inhibition and leads to apoptotic melanoma cell death. **A** Immunoblotting of protein extracts from 1205Lu cells cultivated on MAF-derived ECM treated with vehicle or 5 µM Vemurafenib and/or 7 µM Imatinib, for the indicated time using antibodies against P-ERK1/2, P-Rb, Survivin, cleaved Caspase 3, RelB, P-p65 and HSP60 as loading control. **B** Immunoblot analysis of protein extracts from siCTRL- or siDDR1/2-transfected SKMEL5 cells plated on FRC-derived ECM in the presence of 2 µM Vemurafenib and 0.01 µM Trametinib for 96 h using antibodies against DDR1, DDR2, P-ERK1/2, E2F1, Survivin, cleaved Caspase 3, RelB, p100/p52 and HSP60 as loading control were used. **C** Illustration of the non-canonical p52/RelB NFκB2 pathway and inhibitors used in the study. **D** Immunoblot analysis of protein extracts from 1205Lu cells cultivated on MAF-derived ECM for 96 h in the presence of Vemurafenib and/or a pan-IKK inhibitor (IKKi, BMS-345541 3 µM) using antibodies against P-ERK1/2, P-Rb, Survivin, RelB, P-p65 and HSP60 as loading control. **E** Immunoblotting of protein extracts obtained from SKMEL5 cells that were plated on FRC-derived ECM and treated with 5 µM Vemurafenib in combination or not with the 5 µM DDR1/2 inhibitor DDR1-IN-1 or 10 µM NIK inhibitor (NIKi) for 96 h. Antibodies against P-ERK1/2, P-Rb, Survivin, cleaved Caspase 3, RelB, p100/p52 and HSP60 as loading control were used.

**Figure 9.**
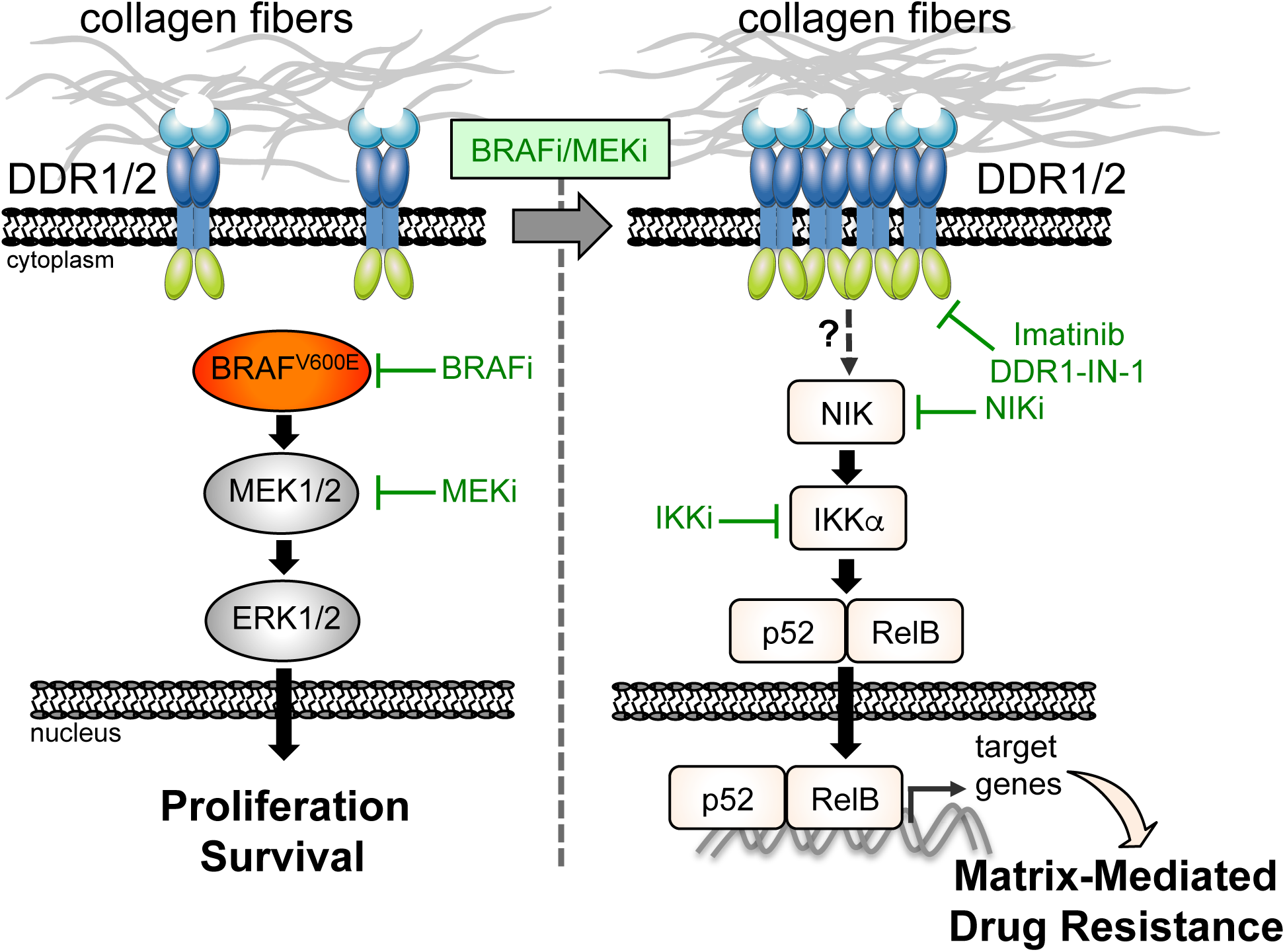
Model of DDR1/DDR2-dependent matrix-mediated drug resistance to MAPK-targeting therapies in melanoma. BRAF-mutated melanoma cells adapt to BRAF/MEK inhibition by turning on a drug tolerance pathway that is initiated by collagen-rich environments interacting with cancer cell DDR1/2. Clustered DDR1/2 activates a non-canonical NFκB2 (p52/RelB) resistance pathway that is therapeutically targetable with clinically approved compounds such as Imatinib or with pre-clinically tested NIK inhibitors.

## Discussion

Preventing melanoma resistance and relapse to therapies targeting the BRAF oncogenic pathway remains a significant challenge in successful disease management. Emerging evidence suggests that stromal components of the tumor microenvironment including the ECM play a key role in establishing resistant niches by allowing melanoma cells to rapidly adapt and tolerate therapeutic drugs before mutation-driven resistance mechanisms are acquired (Meads et al., 2009, Rambow et al., 2019, Smith et al., 2016). We describe here a novel mechanism of adaptation and tolerance to oncogenic BRAF pathway inhibition involving dynamic interaction of melanoma cells with the ECM. ECM molecular composition, fiber orientation and physical characteristics are profoundly altered in the vast majority of solid tumors (Pickup et al., 2014). By comparing cell-derived matrices produced by normal fibroblast and MAFs, we find that biochemical and mechanical properties of experimentally derived matrices differ remarkably depending on fibroblast origin. MAF-derived ECMs exhibit a high level of fiber organization and increased stiffness compared with ECMs generated by normal dermal fibroblasts. This is in agreement with that has been demonstrated in other studies (Gopal et al., 2017, Kaukonen et al., 2017). Interestingly, we disclose that FRCs, which are resident fibroblasts of the lymph node harbor some phenotypic and functional properties of MAFs. Similar to MAFs, FRCs produce and remodel a stiff ECM enriched in fibrillar collagens. Consistent with this, MAF- and FRC-derived matrices display higher drug protective efficacy to melanoma cells than ECM generated from dermal fibroblasts. However, it is important to note that HDF-derived ECM can confer protection against BRAF inhibition in contrast to classic 2D culture conditions, indicating that MM-DR is dependent of the structural organization of the ECM. Importantly, MM-DR was described with several BRAF mutant melanoma cells, regardless their transcriptional phenotypic signature. FRCs play key role in lymph node immune homeostasis (Fletcher et al., 2015). Our *in vitro* data further indicate that similar to MAFs they have the ability to build a protective ECM niche, which might have potential implications for tumor cell behavior and drug tolerance within the lymphatic metastatic site.

Functionally, we demonstrate that collagen receptors DDR1 and DDR2 mediate MM-DR to oncogenic BRAF targeted therapies through a pro-survival NIK-IKKα-NFκB2 pathway. DDR1 and DDR2 are unique among the receptor tyrosine kinases family as they represent non-integrin collagen receptors (Leitinger, 2014, Shrivastava et al., 1997, Vogel et al., 1997). Their role during embryonic development, wound healing and fibrosis is well described (Cario, 2018, Leitinger, 2014). On the contrary, their function during cancer development is less clear as they can act either as tumor-promoters or tumor-suppressors, according to the tissue of origin (Valiathan et al., 2012). Moreover, the functional implication of DDR1 or DDR2 kinase activity in mediating sensitivity to anti-cancer therapies remains poorly defined. DDR1 and DDR2 are both expressed at different levels in skin (Cario, 2018), particularly in the epidermis from which cutaneous melanomas originate, following the malignant transformation of melanocytes (Shain & Bastian, 2016b). Overlapping functions have been attributed to DDR1 and DDR2 in melanoma development with regards to cancer cell growth and invasiveness. A recent study correlated high DDR1 expression in melanoma lesions with poor prognosis and showed that DDR1 controls melanoma cell invasion and survival *in vitro* and in a xenograft melanoma model (Reger de Moura et al., 2019). Other studies have reported that DDR2 depletion in melanoma cell lines reduces their invasive and metastatic abilities (Badiola et al., 2011, Poudel et al., 2015). In line with these observations, analysis of clinical samples representative of melanoma progression displays augmented DDR1 and DDR2 during malignant transition from benign melanocytic lesions to metastatic melanoma. However, examination of melanoma TCGA datasets also indicates a trend towards mutual exclusivity according to high levels of DDR1 or DDR2 mRNA expression, suggesting that their individual function may be coupled to different melanoma phenotypic states. Consistently, analysis of DDRs expression in melanoma cells indicate that high levels of DDR2, but not DDR1, is associated with the AXL^high^ invasive signature also found in therapy-resistant melanoma cells (Muller et al., 2014, Nazarian et al., 2010, Rathore et al., 2019). Whether DDR1 and/or DDR2 expression and signaling contribute to the late phase of acquired resistance remains an open question.

Tumor cell phenotypic switching, which leads to melanoma intra-tumor heterogeneity is currently viewed as a major source of therapeutic escape and relapse (Rambow et al., 2019). In this context, collagen abundance has recently been identified as an important contributor to melanoma phenotype switching through lineage-specific microenvironment sensing (Miskolczi et al., 2018). Moreover, consistent with our observations on BRAFi-treated melanoma tumors, MAPK pathway blockade was shown to increase collagen synthesis *in vitro* (Jenkins et al., 2015, Titz et al., 2016) and collagen deposition *in vivo,* in mutant BRAF cells (Jenkins et al., 2015). We provide evidence for a role of DDR1 and DDR2 during the adaptive phase of tolerance to targeted therapies through MM-DR. Indeed it is tempting to propose that the collagen receptors DDRs represent a major component of the microenvironment-sensing arsenal, driving tumor cell phenotypic plasticity and drug adaptation. Interestingly, β1 integrin, another ECM receptor that plays a major role in EM-DR (Seguin et al., 2015), has also been linked to adaptive responses to BRAF inhibition via fibronectin-mediated activation of FAK, leading to MAPK pathway reactivation (Fedorenko et al., 2015, Hirata et al., 2015). Contrary to β1 integrin, engagement of DDRs by fibrillar collagens mediates drug tolerance in the absence of the reactivation of the MAPK pathway. This suggests that DDR1 and DDR2 participate in the early response to BRAF inhibition through a pathway distinct from β1 integrin engagement.

Although DDR1 and DDR2 display different affinities for collagens, DDR1 is activated by both fibrillar and non-fibrillar collagens whereas DDR2 is activated only by fibrillar collagens (Valiathan et al., 2012). The ECM produced by different types of fibroblasts was found enriched in various matrisome components, including fibrillar and non-fibrillar collagens that activate DDR1/2 clustering into linear membrane structures in melanoma cells, resembling those described in other studies (Agarwal et al., 2007, Yeung et al., 2019). Importantly, receptor clustering on collagen-rich matrices is enhanced upon melanoma cell treatment with BRAFi, thus making the involvement of this process during the early phase of drug adaptation very probable. The exact mechanism underlying the induction of DDR1/2 clustering following MAPK pathway inhibition is however currently unknown. Intriguingly, melanoma cells with acquired resistance to BRAFi exhibit increased actomyosin tension involved in YAP-mediated drug resistance (Kim et al., 2016). It is therefore possible that similarly to acquired resistance the adaptive response of melanoma cells to BRAF inhibition implicates a mechanical remodeling of the actomyosin cytoskeleton. This would result in the assembly of DDR1/2 into supramolecular membrane structures associated with collagen fibrils. Supporting this hypothesis, DDR1 clustering has been shown to involve interactions with collagen and myosin IIA in mice periodontal tissues (Coelho et al., 2017).

As mentioned above, we and others (Jenkins et al., 2015, Titz et al., 2016) have shown that increased collagen deposition is an early response to BRAF inhibition. Our results further revealed that BRAFi treatment increased collagen fiber deposition (Fig 7) and tumor stiffening *in vivo* (data not shown). This supports the notion that DDR1/2 could mediate mechanical signaling from MAF- or FRC-derived stiff ECM, influencing melanoma drug resistance through the creation of a protective niche. Together, these observations support a model in which BRAF targeted drugs fuel a self-feeding mechanism involving collagen-bound DDR1/2 signaling platforms, responsible for collagen network compaction and drug tolerance. We show that DDR1/2 knockdown or the inhibition of their catalytic activity impairs drug tolerance. It would thus be interesting to examine how genetic or pharmacological inhibition of the two receptors affects the assembly of their signaling platforms on melanoma cells. Another important issue revealed by our *in vivo* model of melanoma drug response is the possible implication of stromal DDR1/2 in the therapeutic response. Consistent with this a major role of CAF-derived DDR2 for the organization of collagen fibers and breast cancer metastasis has been reported (Corsa et al., 2016). DDR1/2 signaling might therefore influence both cancer and stromal cells during tumor adaptation to BRAF inhibition.

To gain insight into DDR1/2-related signaling pathways, we performed a biochemical screen to identify tyrosine kinases or kinase substrates potentially involved in the MM-DR process. Unfortunately, this approach failed probably due to the lack of regulators of the NFκB pathway in the phospho-antibody array employed. Subsequent biochemical studies revealed that fibroblast-derived matrices enhanced the activation of the non-canonical-NFκB pathway that accounted for most of melanoma cell tolerance to BRAF inhibition. The NFκB2/p52-RelB pathway represents a major alternative route of NFκB signaling (Taniguchi & Karin, 2018), that has been involved in cell adhesion-mediated drug resistance (CAM-DR) in myeloma (Landowski et al., 2003) and in drug resistance in prostate cancer (Nadiminty, Tummala et al., 2013). In melanoma, the NFκB2 pathway is upregulated compared to melanocytes and prevents melanoma senescence through increase expression of the Polycomb-group protein EZH2 (De Donatis et al., 2016). Here we show that suppression of DDR1/2 signaling impaired MM-DR by reducing the expression of NFκB2/p52. Interestingly, the siRNA approach indicated that DDR1 and DDR2 compensate for one another, as we observed that both collagen receptors must be knock-downed to fully prevent MM-DR. Consistently, DDR1/2 inhibition by the clinically approved TKI Imatinib improved the action of BRAFi on melanoma treatment by delaying tumor relapse, normalizing collagen network and increasing mice survival. Thus, this combination strategy could be effective in a clinical application as it allows the suppression of the stromal fibrotic reaction induced by oncogenic BRAF pathway inhibition and prevents tumor relapse. In addition, we observe that the inhibition of NIK or IKK, two upstream activators of the non-canonical NFkB pathway (Taniguchi & Karin, 2018), also abrogated ECM-mediated resistance to BRAFi. How DDR1/2 connect to the NIK-IKK-NFκB2/p52 pathway is currently unknown. However, it is likely that the DDR1/2 signaling platforms that were identified are implicated. The nature of the NFκB2/p52 target genes that mediate MM-DR in melanoma is also under investigation.

Finally, our work adds to the emerging notion that DDR1 and DDR2 are becoming attractive targets in anti-cancer therapies. Inhibiting DDR1 or DDR2 with Nilotinib (Jeitany et al., 2018), Dasatinib (Hammerman et al., 2011, von Massenhausen et al., 2016), or other non-approved inhibitors (Ambrogio et al., 2016, Grither & Longmore, 2018) has been shown to decrease tumorigenicity, invasion or metastasis of several types of carcinomas (Corsa et al., 2016) Importantly, a recent study in melanoma and other solid tumors reported that targeting DDR2 with Dasatinib enhances tumor response to anti-PD-1 immunotherapy (Tu et al., 2019). Together this later study and our study point to the critical role of these collagen receptors in regulating the immune and mesenchyme tumor stroma in melanoma.

In summary, our findings reveal that DDR1/2-mediated MM-DR favors the persistence of drug-tolerant tumor cells during the initial phase of adaptation to MAPK-targeting therapies. We also provide evidence that targeting MM-DR with clinically approved DDR inhibitors may represent an attractive salvage strategy to overcome resistance to oncogenic BRAF inhibition. This combinatorial approach may thus be beneficial for melanoma patients on targeted therapies.

## Materials and Methods

### Melanoma cells, reagents and antibodies

Melanoma cell lines were obtained as previously described (Didier et al., 2018, Rathore et al., 2019, Tichet et al., 2015). Isogenic pairs of Vemurafenib-sensitive (P) and resistant (R) cells (M229, M238, M249) were provided by R.S. Lo (Nazarian et al., 2010). Short-term cultures of patient melanoma cells MM034, MM074, MM029, MM099 were kindly provided by J.-C. Marine and were described elsewhere (Verfaillie, Imrichova et al., 2015). Melanoma cells were cultured in Dulbecco’s modified Eagle medium (DMEM) supplemented with 7% fetal bovine serum (FBS) (HyClone) and 1% penicillin/streptomycin solution. Cell lines were used within 6 months between resuscitation and experimentation. To guarantee cell line authenticity, cells were expanded and frozen at low passage after their receipt from original stocks, used for a limited number of passages after thawing and routinely tested for the expression of melanocyte lineage proteins such as MITF. All cell lines were routinely tested for the absence of mycoplasma by PCR. For live imaging and red nuclear labeling 501Mel and 1205Lu cells were transduced with NucLight Red lentivirus reagent (Essen Bioscience) and selected with puromycin (1 μg/ml, Sigma Aldrich).

Culture reagents were purchased from Thermo Fisher Scientific. BRAFi (PLX4032, Vemurafenib), MEKi (GSK1120212, Trametinib), dual RAFi/MEKi (RO5126766), DDR1-IN-1, Nilotinib, and IKK inhibitor BMS345541 were from Selleckchem. Imatinib mesylate was from Enzo Lifescience. NIK inhibitor (NIKi) was described before (Mondragon et al., 2019). FAK inhibitor PF573228 was from Tocris Bioscience. An equal amount of DMSO was used as vehicle control. Collagen I was from Thermo Fisher Scientific. All other reagents were obtained from Sigma-Aldrich unless stated otherwise. β1 Integrin blocking antibody (clone AIIB2) and control isotype were from Merck Millipore. Information on antibodies used in this study is provided in Supplementary Table 1.

### Isolation and culture of primary fibroblasts and MAFs

Human dermal fibroblasts (HDFs) were isolated and maintained as described previously (Robert, Gaggioli et al., 2006). Human lymphatic fibroblasts (FRC#1 and #2) were purchased from ScienCell Research Laboratories. Metastatic melanoma clinical specimens were obtained with written informed consent from each patient and the study was approved by the hospital ethic committee (Nice Hospital Center and University Côte d’Azur). Briefly, the sample was cut into small pieces and digested using collagenase/dispase. Following filtration of the large debris, the solution was serial centrifuged and the final pellet was re-suspended in DMEM supplemented with 10% FBS and seeded in a tissue culture dish. After 30 min, the fibroblasts adhered to the dish whilst other cellular types remained in suspension. Fibroblasts were cultured in fibroblast medium (ScienCell) supplemented with 10% FBS and 1% fibroblast growth supplement (FGS) plus 1% penicillin/streptomycin solution. All experiments were performed with fibroblasts until passage 10. Verification of the identity of MAFs was determined by flow cytometry using classical markers.

### Fibroblast-derived 3D ECM and MM-DR assay

3D de-cellularized ECMs were generated as previously described (Beacham et al., 2007). Briefly, gelatin-coated tissue culture dishes were seeded with fibroblasts and cultured for 8 days in complete medium, supplemented with 50 μg/ml ascorbic acid every 48 h. Cells were then washed with PBS and matrices were de-cellularized using a pre-warmed extraction buffer for 2 minutes (PBS 0.5% Triton X-100, 20 mM NH_4_OH). Matrices were then gently washed several times with PBS.

For matrix-mediated drug-resistance (MM-DR) assays, melanoma cells were seeded ontop of the de-cellularized 3D matrices for 48 h at 37°C in 5% CO_2_, and cultured in complete medium for a further 48 to 96h in the presence of the various inhibitors, as indicated in the figure legends. Cells were detached and fixed in 80% ethanol or lysed in lysis buffer for cell cycle and immunoblot analysis, respectively.

#### Matrix remodeling assay

5 × 10^4^ fibroblasts were embedded in 100 µl of collagen I/Matrigel and seeded in a glass bottom 96-well plate (MatTek) and maintained in DMEM supplemented with 10% FBS. Gel contraction was monitored until day 6. The gel area was measured using ImageJ software and the contraction was calculated using the formula 100 × (well diameter−gel diameter)/well diameter as previously described (Albrengues et al., 2014).

### RNAi studies

Non-targeting control, DDR1#1 (VHS50139), DDR1#2 (HSS187878), DDR2#1 (HSS107350), DDR2#2 (HSS107352) and FAK (PTK2) siRNA duplexes were purchased from Thermo Fisher Scientific. Transfection of siRNA was carried out using Lipofectamine RNAi MAX (Thermo Fisher Scientific), at a final concentration of 50 nM. Unless stated otherwise, cells were assayed at 2 or 4 days post transfection.

### Immunoblot and immunoprecipitation

Whole-cell lysates were prepared using lysis buffer containing 50 mM Hepes, 150 mM NaCl, 1.5 mM MgCl_2_, 1 mM EDTA, 10% Glycerol, 1% Triton X-100 supplemented with protease and phosphatase inhibitors (Pierce) and briefly sonicated. Proteins were separated by SDS-PAGE and were transferred onto PVDF membranes (GE Healthcare Life sciences) for immunoblot analysis. Membranes were incubated with the primary antibody overnight, washed and then incubated with the peroxidase-conjugated secondary antibody. Antibodies and working dilution used are listed in Supplementary Table 1. Blots were developed with a chemiluminescence system (GE Healthcare Life Sciences).

For IP assays, cells treated with 10 μg/ml of collagen I for 18 h with the indicated inhibitors were lysed as described above, then incubated in lysis buffer containing Protein G Sepharose beads (Merck) and an antibody directed against DDR2 overnight at 4°C with rocking. Beads were then washed three times with 20 mM Hepes, 150 mM NaCl, 10% Glycerol, 0.1% Triton X-100 supplemented with protease and phosphatase inhibitors. IP products were separated by SDS-PAGE and subjected to immunoblot analysis.

### Proliferation assay

For real time analysis of cell growth using the IncuCyte^TM^ ZOOM imaging system (Essen Bioscience), cells stably expressing the nuclear fluorescent label red NucLight reagent were plated in quadruplicate in complete medium (5 x 10^3^ cells/well for 501MEL and 15 x 10^3^ cells/well for 1205Lu cells) in coated (fibroblast-derived ECM or Collagen type I) or uncoated wells in a 12 well plates. Phase contrast images were taken every hour over a 5-day period. Cell proliferation was quantified by counting the number of fluorescent nuclei over time to give cell growth rate. Growth curves were generated using the IncuCyte^TM^ cell proliferation assay software. Alternatively, cell proliferation was analyzed by counting cells with Hoechst-stained nuclei.

### Flow cytometry

Cell cycle profiles were determined by flow cytometry analysis of propidium iodide (PI)-stained cells as previously described (Didier et al., 2018). Melanoma cells cultured on top of fibroblast-derived matrices and treated with the indicated drugs for 48 h were washed, fixed in 80% ethanol and incubated at −20°C for 24 h. Cells were then stained for 20 min at 37°C in buffer containing 40 μg/ml PI and 20 μg/ml ribonuclease A. Cell cycle profiles were collected on a FACSCanto II (Becton Dickinson). Cell death was evaluated following staining with AnnexinV/PI (eBioscience) and analyzed by flow cytometry as described (Didier et al., 2018).

### Immunofluorescence and microscopy

Cells were grown on type I collagen-coated glass coverslips (0.14 mg/ml) or fibroblast derived ECM-coated glass coverslips. After Vemurafenib treatment, cells were rinsed with PBS, fixed in 4% paraformaldehyde (PFA), and incubated in PBS containing 10% normal goat serum (Cell Signaling) for 1 h, then incubated overnight with the indicated primary antibodies diluted in PBS containing 2% normal goat serum. Following incubation with Alexa Fluor-conjugated secondary antibodies, coverslips were mounted in ProLong antifade mounting reagent (Thermo Fisher Scientific). F-actin was stained with Alexa Fluor 488 phalloidin (1:100; Thermo Fisher Scientific). Nuclei were stained with DAPI. Images were captured using a wide field (Leica DM5500B, at x 63 magnification) or Zeiss LSM 510 META laser scanning confocal microscopes. Cell area and globular *versus* linear clusters of cells (n > 50) were determined and quantified using the ImageJ software.

Following fixation and incubation with primary antibodies and Alexa Fluor-conjugated secondary antibodies, de-cellularized matrices on coverslips were mounted in ProLong antifade reagent. Images were captured using a wide field microscope (Leica DM5500B, at x 40 magnification). The orientation of fibronectin and collagen fibers was assessed in the immunofluorescence images using ImageJ software. Data were plotted as frequency of distribution.

Cleaved caspase 3 and Ki67 staining was assessed on 5 μm frozen sections of cell-derived melanoma xenografts. Samples were fixed for 30 min with 3% PFA in PBS, rehydrated for 10 min in Tris 0.1 M, permeabilized in Tris 0.1 M + 1% Triton for 1 h, and then blocked for 1 h in Tris 0.1M containing 1% BSA, 0.3% Triton, anti-CD16/CD32 (FC Block, 1/100). Antibodies were diluted in Tris 0.1M containing 1% BSA, 0.3% Triton and incubated overnight at 4°C. After washes with Tris 0.1M, bound antibodies were detected using Alexa Fluor 488 or Alexa Fluor 594-conjugated secondary antibody. Images were captured using a wide field microscope (Leica DM5500B, at x 40 magnification).

### Mass spectrometry analysis

Proteomic analysis of de-cellularized matrices was performed as described in Gopal *et al*. (Gopal et al., 2017). Briefly, ECM proteins were solubilized in urea, reduced and alkylated and proteins were digested with first PNGase F (New England BioLabs), endoproteinase Lys-C (Promega) and high-sequencing-grade trypsin. Each sample was reconstituted in 0.1% trifluoroacetic acid (TFA) 2% acetonitrile and analyzed by liquid chromatography (LC)–tandem mass spectrometry (MS/MS) in an LTQ-Orbitrap-Velos (Thermo Electron, Bremen, Germany) online with a nanoLC Ultimate 3000 chromatography system (Dionex). For protein identification and estimation of abundance, the acquired raw LC Orbitrap MS data were processed using the MASCOT search engine (version 2.4.1). Spectra were searched against a SwissProt Human database. The protein abundance was calculated using the iBAQ score and represented as molar percent.

### Phospho-kinase profiling

Phospho-kinase screening was performed using a phospho-kinase array (Proteome Profiler Human Phospho-Kinase Array #ARY003B; R&D Systems) according to the manufacturer’s instructions. After cell extraction, 600μg of protein was added per sample. Array spots were analyzed using ImageJ software.

### Atomic force microscopy

Mechanical properties of fibroblast-derived matrices were analyzed by AFM using a Bioscope Catalyst operating in Point and Shoot (Bruker Nano Surfaces), coupled with an inverted optical microscope (Leica DMI6000B, Leica Microsystems Ltd.). The apparent Young’s Modulus (E_app_) was measured on unfixed ECM using a Borosilicate Glass spherical tip (5 μm of diameter) mounted on a cantilever with a nominal spring constant of 0.06 N/m (Novascan Technologies). The force-distance curves were collected using a velocity of 2 μm/s, in relative trigger mode and by setting the trigger threshold to 1 nN. E_app_ values were represented as a boxplot using GraphPad Prism (GraphPad software).

### Cell line-derived xenograft (CDX) tumor models

Mouse experiments were carried out in accordance with the Institutional Animal Care and the local ethical committee (CIEPAL-Azur agreement NCE/2018-483). 1 × 10^6^ 1205Lu melanoma cells were subcutaneously implanted into both flanks of 6 week old female athymic nude nu/nu mice (Janvier, France). The tumor was measured by caliper and the volume was calculated using the formula: V = tumor width x tumor length2 x 0.5. When the tumor reached 75 mm^3^, mice were randomly grouped into control and test groups. Vemurafenib (35 mg/kg) and Imatinib mesylate (75 mg/kg) were delivered (alone or in combination) intraperitoneally three times per week. Mice in the control group were treated with vehicle alone. Mice were treated for 30 days and followed for up to 50 days or until tumors reached a pre-defined volume (1000 mm^3^). After animal sacrifice, tumors were dissected, weighed, and snap frozen in liquid nitrogen in optimal cutting temperature compound (OCT; Tissue-Tek) (Gentaur) for immunofluorescence analysis or formalin fixed and paraffin embedded for picrosirius red staining or SHG analysis.

### Fibrillar collagen imaging

Collagen in de-cellularized matrices or in paraffin-embedded melanoma tissues was stained with picrosirius red using standard protocols. Tumor sections were analyzed by polarized light microscopy as described (Rich & Whittaker, 2005). Images were acquired under polarized illumination using a light transmission microscope (Zeiss PALM, at 5 x magnification). Fiber thickness was analyzed by the change in polarization color. Birefringence hue and amount were quantified as a percent of total tissue area using ImageJ software.

Second harmonic generation (SHG) imaging of paraffin-embedded melanoma tissues was recorded on a Zeiss 510 NLO microscope (Carl Zeiss Microscopy) with Mai Tai HP DeepSee (Newport Corporation) and 360-440 nm band pass filter.

### Tissue microarray analysis

Immunohistochemistry analysis of DDR1 and DDR2 expression was assessed on TMA sections (US Biomax) using VECTASTAIN Elite ABC Kit (Vector Laboratories) with the DAB reagent (Vector Laboratories) and counterstained with haematoxylin (Fisher Scientific), following the manufacturer’s instructions. Images were captured on a bright field microscope (Nikon, at 20 x magnification).

### Analysis of gene expression from public databases

Publicly available gene expression data sets of human melanoma samples were used to analyze *DDR1 and DDR2* levels in the Mannheim (GSE4843), Philadelphia (GSE4841), and Zurich (GSE4840) cohorts. Proliferative and invasive melanoma subgroups were defined as previously described (Rathore et al., 2019, Widmer et al., 2012). Survival data from the skin melanoma TCGA database were retrieved using cBioPortal (cbioportal.org).

### Statistical analysis

Unless otherwise stated, all experiments were repeated at least three times and representative data/images are shown. Statistical data analysis was performed using GraphPad Prism 8 software. The unpaired two-tailed Mann-Whitney test was used for statistical comparisons between two groups. The Kruskal-Wallis test with the indicated post-tests or two-way analysis of variance test with Sidak’s post-test was used to compare three or more groups. Error bars are ± SEM.

## Acknowledgments

We thank Roger Lo for M229P/R, M238P/R and M249P/R melanoma cells and J.C. Marine for short-term cultured melanoma cells. We acknowledge Maeva Gesson and Marie Irondelle from the C3M imaging facility for their helpful advice. We thank the C3M animal facility and Cédric Matthews for the SHG microscopy analysis (IBDM imaging platform, AMU-Marseille). We also thank Ana Popovic for critical reading of the manuscript. This work was supported by funds from the: Institut National de la Santé et de la Recherche Médicale (Inserm), Ligue Contre le Cancer, Institut National du Cancer (INCA_12673), ITMO Cancer Aviesan (Alliance Nationale pour les Sciences de la Vie et de la Santé, National Alliance for Life Science and Health) within the framework of the Cancer Plan, and the French Government (National Research Agency, ANR) through the ‘’Investments for the Future’’ LABEX SIGNALIFE: program reference # ANR-11-LABX-0028-01. We also thank the platform PiCSL-FBI (IBDM, AMU-Marseille) a member of France BioImaging supported by the ‘’Investments for the Future’’ (ANR-10-INBS-04). The financial contribution of the Conseil général 06, Canceropôle PACA and Region Provence Alpes Côtes d’Azur to the C3M is also acknowledged. I.B. was a recipient of a doctoral fellowship from La Ligue Contre le Cancer.

## Author contributions

S.T.D. and M.D. designed the study, analyzed and interpreted data, ensured financial support, and wrote the paper with inputs from I.B., V.P. and C.A.G. I.B. performed the majority of the experiments and analyzed the data with the help of M.L, A.M., S.D., C.R., V.P. and C.A.G. F.L. performed flow cytometry analyses. S.P. performed AFM analyses. S.A. performed mass spectrometry analyses. C.G. and T.P. contributed with reagents and expertise.

## SUPPLEMENTARY FIGURE LEGENDS

**Supplementary Figure 1.**
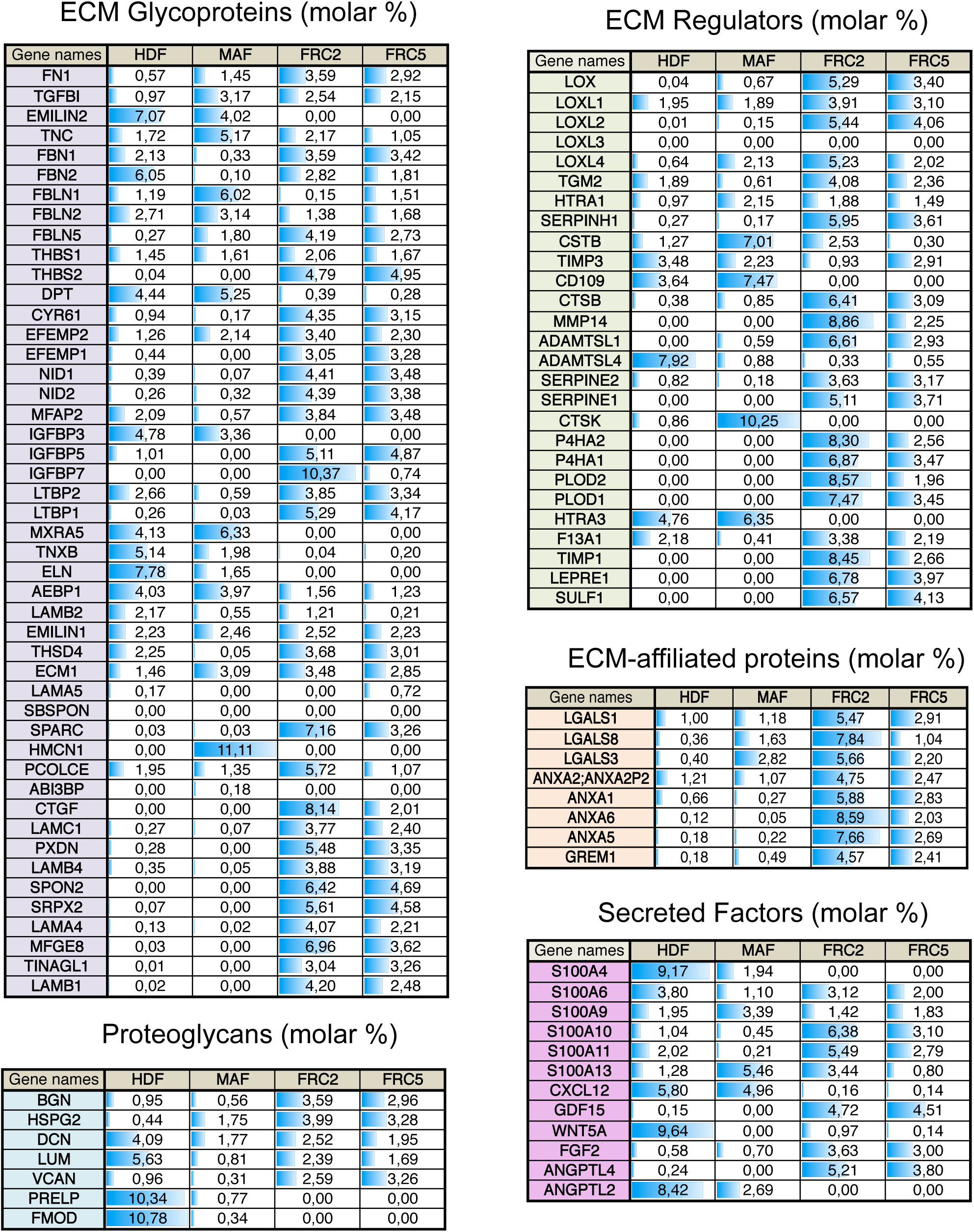
Quantitative mass spectrometry analysis of fibroblast-derived ECM. Complete list of core matrisome and matrisome-associated proteins detected by mass spectrometry of fibroblast-derived ECM produced by HDF, skin MAF or by two lymphatic FRCs (FRC#2 and FRC#5). The table provides the Mascot results and the molar % (iBAQ score) for identified proteins at 1% FDR. Identified proteins were matched with the human matrisome database (Naba A, Ding H, Whittaker CA, Hynes RO. http://matrisomeproject.mit.edu) to retrieve the division and the category. The blue histogram and the values correspond to the molar % calculated for the entire dataset or for the different matrisome categories.

**Supplementary Figure 2.**
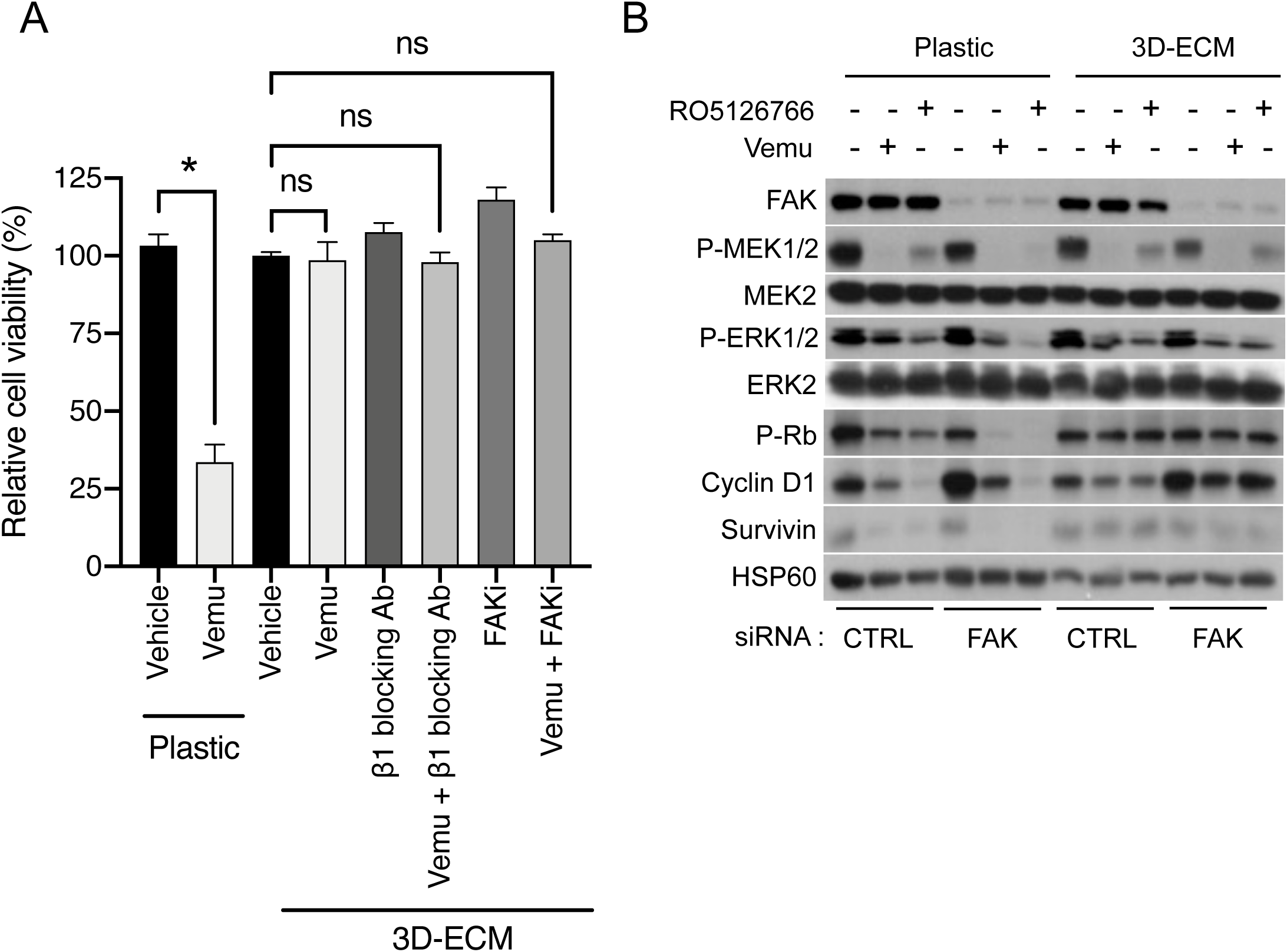
Blocking β1 integrin/FAK pathway does not inhibit ECM-mediated resistance to BRAF inhibition. **A** Quantification of 501MEL cell proliferation following 48 h of culture on plastic or FRC-derived ECM prior to treatment with 2 µM Vemurafenib in the presence or not of β1 integrin blocking antibodies (10 µg/ml) or FAK inhibitor (5 µM) for 7 h. ns, non significant, *P<0.05, Kruskal-Wallis test. **B** Immunoblotting of protein extracts from siCTRL- or siFAK-transfected 501MEL cells plated on FRC-derived ECM treated with vehicle or 2 µM Vemurafenib or 1 µM dual RAF/MEK inhibitor RO5126766 for 72 h, using antibodies against FAK, P-MEK1/2, P-ERK1/2, P-Rb, Cyclin D1 or survivin. HSP60, loading control.

**Supplementary Figure 3.**
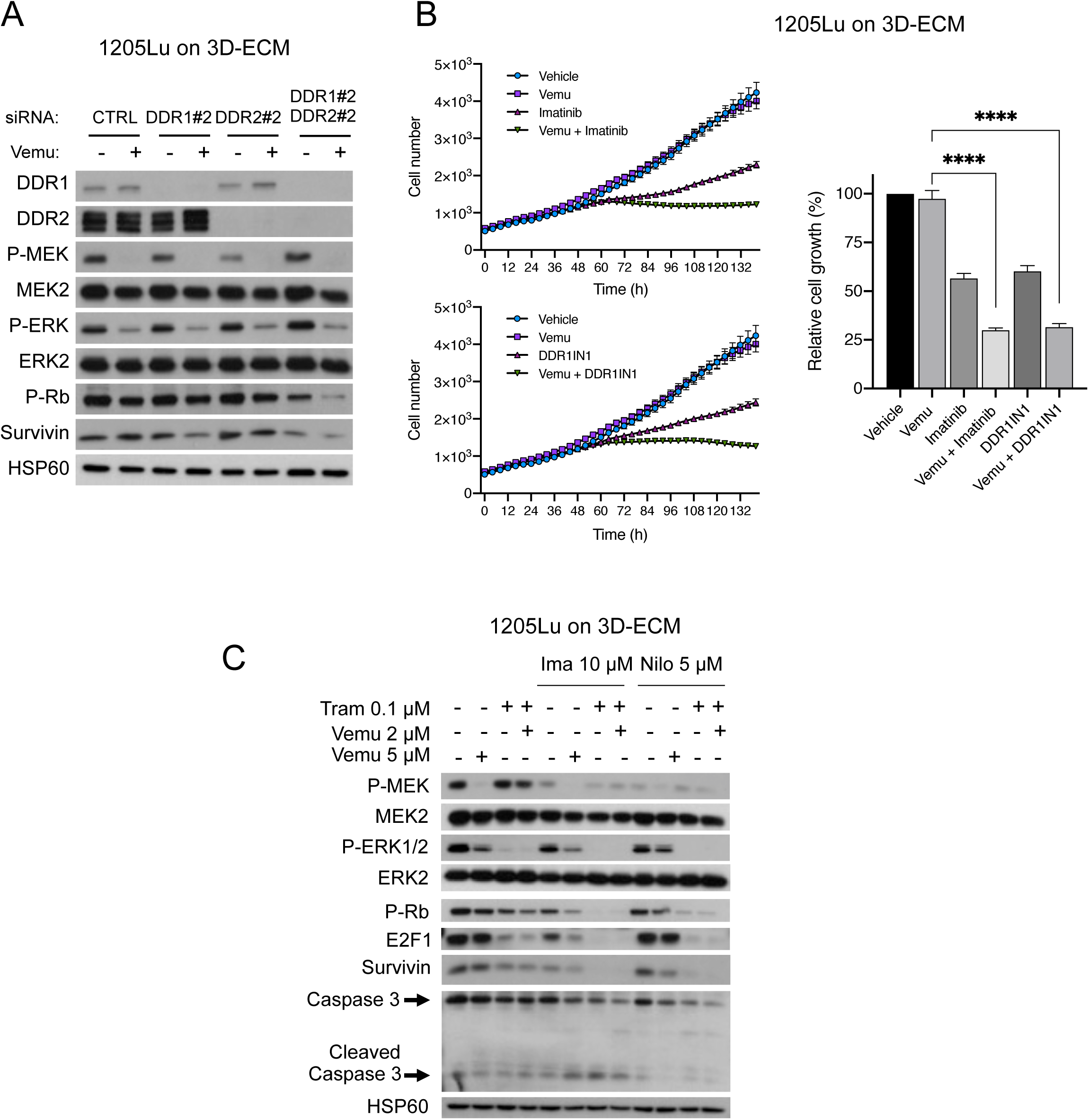
Inhibition of DDR1 and DDR2 by genetic or pharmacological approaches abrogates ECM-mediated resistance to BRAF^V600E^ pathway inhibition. **A** Immunoblotting of protein extracts of siCTRL-, siDDR1#2-, siDDR2#2- or siDDR1#2/siDDR2#2- transfected 1205Lu cells plated on MAF-derived ECM in the presence or not of 5 µM Vemurafenib for 96 h, using antibodies against DDR1, DDR2, P-MEK1/2, P-ERK1/2, P-Rb or survivin. HSP60, loading control (n=2). **B** Time-lapse imaging of proliferation of NucLight-labeled 1205Lu plated for 48 h on FRC-derived ECM prior treatment with 5 µM Vemurafenib in the presence or not of 7 µM Imatinib or 1 µM DDR1-IN-1 for the indicated time. Right bar histograms show quantification of cell proliferation at the experiment end point. ****P<0.001, Kruskal-Wallis test. **C** Immunoblotting of protein extracts from 1205Lu cells cultivated on FRC-derived ECM for 96 h in the presence of 5 µM Vemurafenib, 0.01 µM Trametinib or the combination of 2 µM Vemurafenib and 0.01 µM Trametinib, in the presence or not of 5 µM Imatinib or 10 µM Nilotinib using anti-P-MEK1/2, P-ERK1/2, P-Rb, E2F1, Survivin, or Cleaved Caspase 3 antibodies (n=2). HSP60, loading control.

**Supplementary Figure 4.**
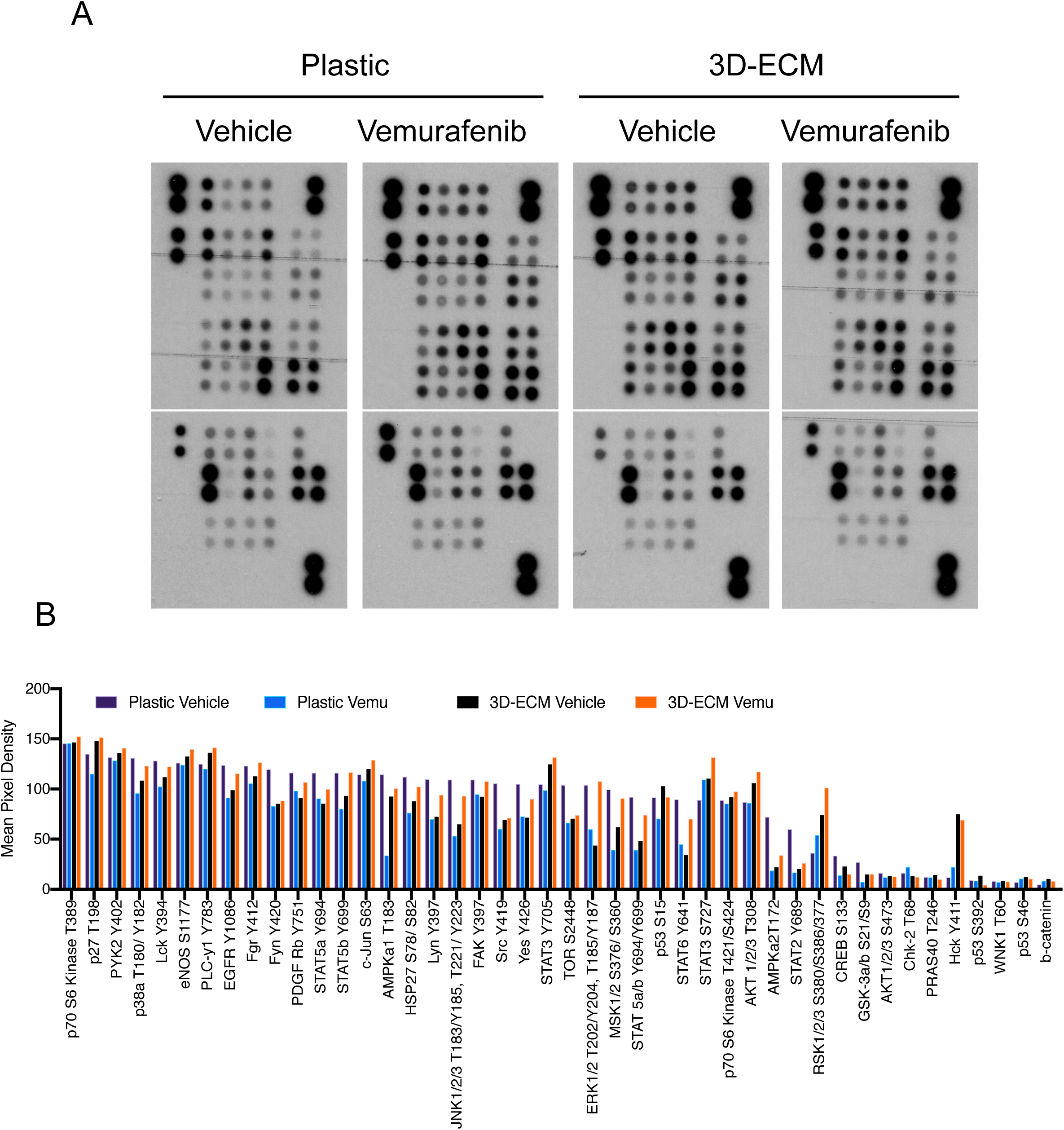
Analysis of the phospho-proteome of BRAFi-treated melanoma cells cultivated on 3D-ECM. **A** Kinases and phosphorylated substrates were detected using an immunoblot array (Proteome Profiler Human phospho-kinase array kit). 501MEL cells were cultured on plastic or 3D-ECM for 48 h prior treatment with 2 µM Vemurafenib for additional 2 days before preparation of cell lysates and immunoblot array analysis following manufacturer’s instructions. **B** Mean pixel density was analyzed using Image J software.

**Supplementary Figure 5.**
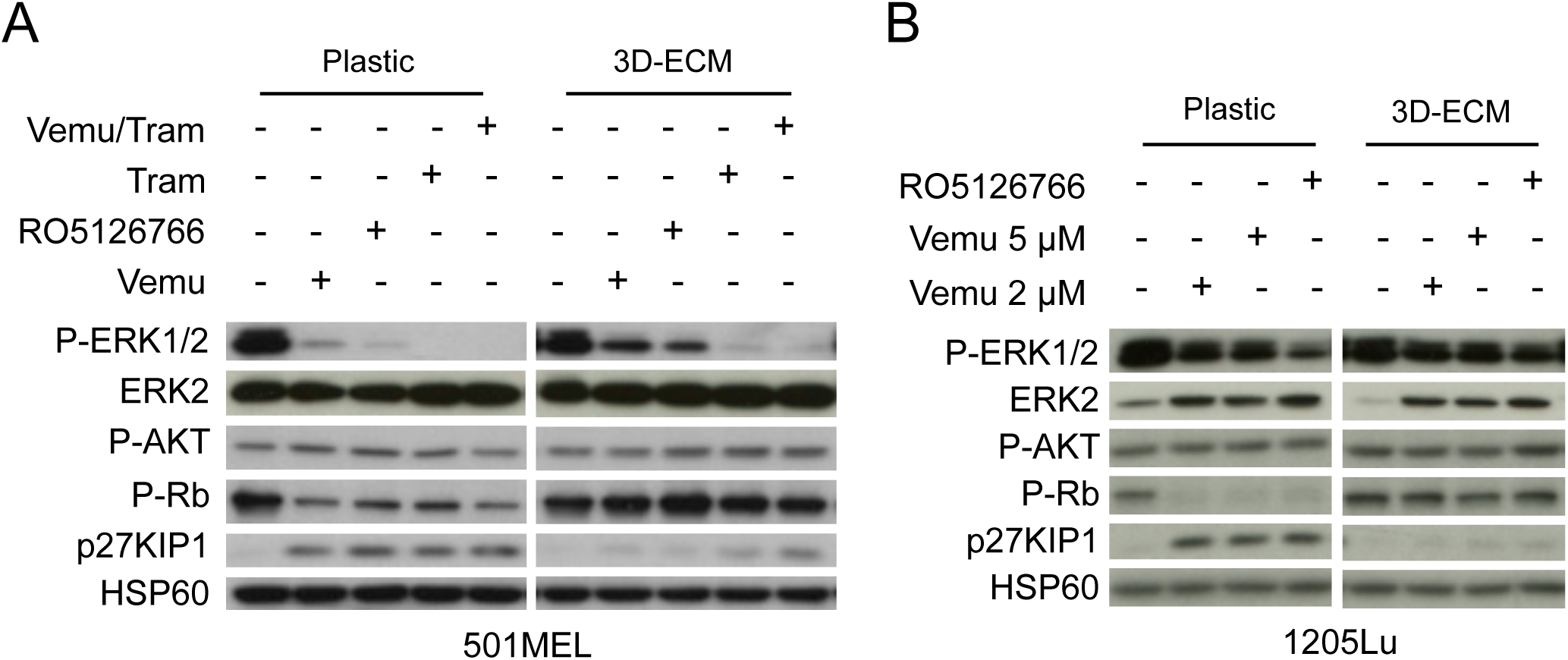
Analysis of phospho-AKT levels in BRAFi/MEKi-treated melanoma cells cultivated on 3D-ECM. **A** Immunoblotting of protein extracts from 501MEL cells plated on plastic or FRC-derived ECM in the presence or not of 2 µM Vemurafenib, 1 µM RO5126766, 0.01 µM Trametinib or the combination 0.01 µM Trametinib/2 µM Vemurafenib for 72 h, using antibodies against P-ERK1/2, P-AKT, P-Rb or p27KIP1. HSP60, loading control. **B** Immunoblotting of protein extracts from 1205Lu cells plated on plastic or FRC-derived ECM in the presence or not of 2 µM or 5 µM Vemurafenib or 1 µM dual RAF/MEKi RO5126766 for 72 h, using antibodies against P-ERK1/2, P-AKT, P-Rb or p27KIP1. HSP60, loading control.

**Supplementary Figure 6.**
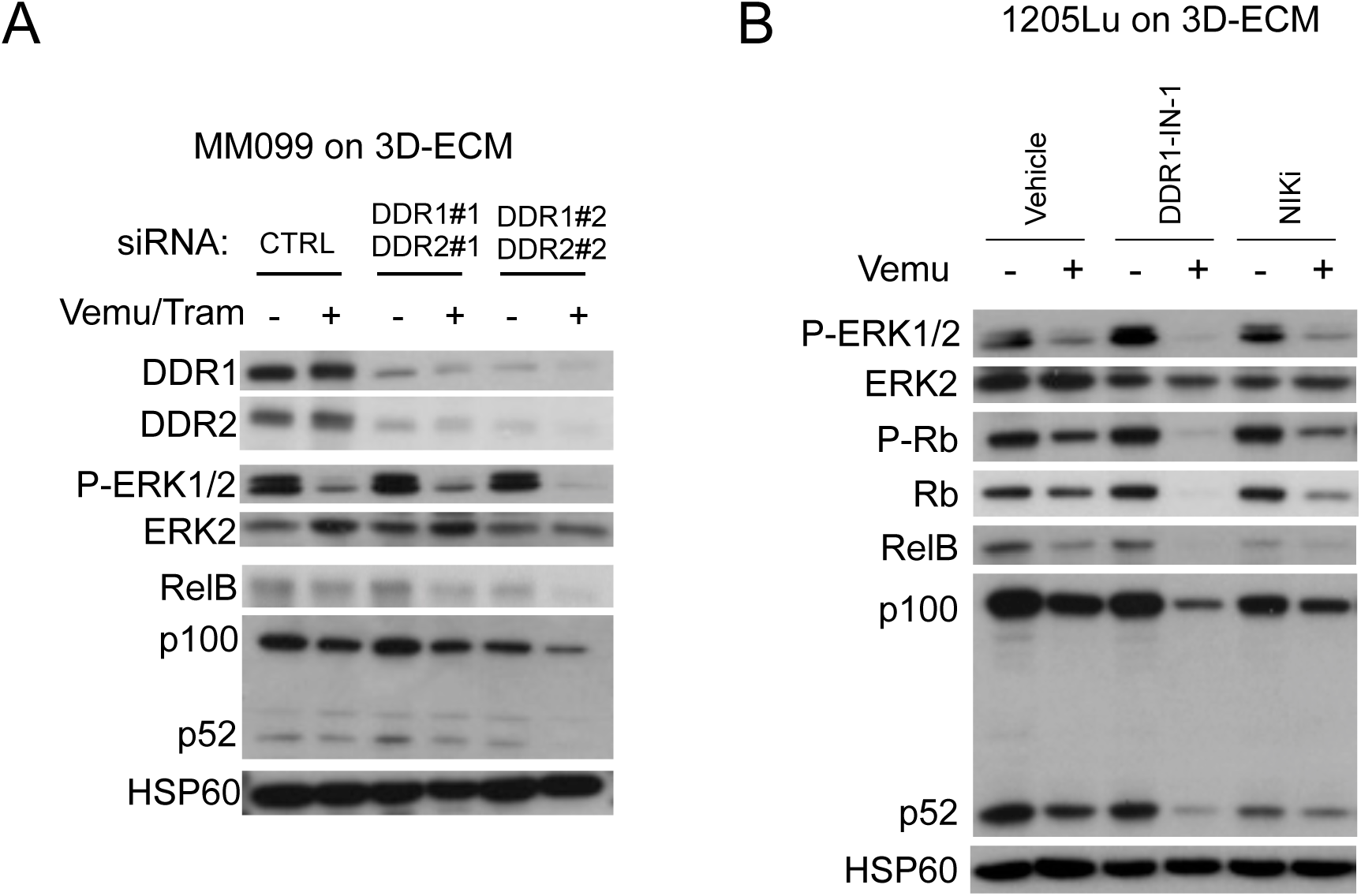
Pharmacological targeting of DDR1/2 or NIK decreases expression of NFκB2/p52 and RelB in ECM-cultivated and BRAFi-treated melanoma cells. **A** Immunoblot analysis of protein extracts from siCTRL- or siDDR1/2-transfected short-term MM099 cells plated on FRC-derived ECM in the presence of 2 µM Vemurafenib and 0.1 µM Trametinib for 96 h. Anti-DDR1, DDR2, P-ERK1/2, RelB, p100/p52 and HSP60 as loading control were used. **B** Immunoblot analysis of protein extracts from 1205Lu cells plated on FRC-derived ECM and treated with or without 5 µM Vemurafenib in the presence or not of DDR1/2 inhibitor (DDR1-IN-1, 5 µM) or NIK inhibitor (NIKi, 10 µM) for 96 h. Antibodies against P-ERK1/2, RelB, p100/p52 and HSP60 as loading control were used.

**Supplementary Table 1:** List of antibodies used in this study.

| **Primary Antibody** | **Company** | **Catalog number** | **Dilutions** |
| --- | --- | --- | --- |
| AXL (C89E7) | Cell Signaling | 8661 | WB 1:1000 |
| BAX | Cell Signaling | 2772 | WB 1:1000 |
| BIM (H-191) | Santa Cruz | sc-11425 | WB 1:1000 |
| Caspase 3 | Cell Signaling | 9662 | WB 1:500 |
| Cleaved Caspase 3 | Cell Signaling | 9661 | IF 1:100 |
| Collagen I | Abcam | ab34710 | IF 1:500 |
| Cyclin D1 (DCS-6) | BD Biosciences | 556470 | WB 1:1000 |
| Cyclin D1 (DCS-6) | BioLegend | 681902 | WB 1:1000 |
| DDR1 (D1G6) XP® | Cell Signaling | 5583 | WB 1:1000 IHC 1:100 |
| DDR2 | Cell Signaling | 12133 | WB 1:1000 |
| DDR2 (3B11E4) | Santa Cruz | sc-81707 | IHC 1:50 |
| E2F1 | Cell Signaling | 3742 | WB 1:1000 |
| ERK2 | Santa Cruz | sc-1647 | WB 1:1000 |
| FAK | Upstate | 05-182 | WB 1:1000 |
| Fibronectin (EP5) | Santa Cruz | sc-8422 | IF 1:200 |
| HSP60 (K19) | Santa Cruz | sc-1722 | WB 1:1000 |
| Ki67 | Abcam | ab16667 | IF 1:250 |
| MEK2 | GeneTex | GTX630542 | WB 1:1000 |
| MITF (C5) | Invitrogen | MA5-14146 | WB 1:1000 |
| NFkB2 p100/p52 (18D10) | Cell Signaling | 3017 | WB 1:1000 |
| p-DDR1 (Tyr792) | Cell Signaling | 11994 | WB 1:1000 IF 1:100 |
| p-DDR2 (Tyr740) | R&D System | MAB25382 | WB 1:1000 IF 1:100 |
| p-ERK1/2 (Thr202/ Tyr204) | Cell Signaling | 9101 | WB 1:1000 |
| p-FAK (Tyr397) | Cell Signaling | 3283 | WB 1:1000 |
| p-MEK1/2 (Ser 221) (166F8) | Cell Signaling | 2338 | WB 1:1000 |
| p-P65 (Ser536) (93H3) | Cell Signaling | 3033 | WB 1:1000 |
| p-PDGFRb (Tyr751) | R&D System | AF1767 | WB 1:500 |
| p-Rb (Ser795) | Cell Signaling | 9301 | WB 1:1000 |
| p27Kip1 (D69C12) | Cell Signaling | 3686 | WB 1:1000 |
| PDGFRb (C82A3) | Cell Signaling | 4564 | WB 1:1000 |
| Rb (4H1) | Cell Signaling | 9309 | WB 1:1000 |
| RelB (C1E4) | Cell Signaling | 4922 | WB 1:1000 |
| SOX10 | Abcam | ab155279 | WB 1:1000 |
| Survivin | Cell Signaling | 2808 | WB 1:1000 |

| **Secondary Antibody** | **Company** | **Catalog number** | **Dilutions** |
| --- | --- | --- | --- |
| Anti-mouse IgG, HRP-linked Antibody | Cell Signaling | 7076 | WB 1:2000 |
| Anti-rabbit IgG, HRP-linked Antibody | Cell Signaling | 7074 | WB 1:2000 |
| Goat anti-Rabbit, Alexa Fluor® 488 | Invitrogen | A11034 | IF 1:200 |
| Goat anti-Rabbit, Alexa Fluor® 594 | Invitrogen | A11012 | IF 1:200 |
| mouse anti-goat IgG-HRP | Santa Cruz | sc-2354 | WB 1:5000 |

